# *C. elegans* nuclear RNAi factor SET-32 deposits the transgenerational heritable histone modification, H3K23me3

**DOI:** 10.1101/2020.02.19.956656

**Authors:** Lianna Schwartz-Orbach, Chenzhen Zhang, Simone Sidoli, Richa Amin, Diljeet Kaur, Anna Zhebrun, Julie Ni, Sam Gu

## Abstract

Nuclear RNAi provides a highly tractable system to study RNA-mediated chromatin changes and epigenetic inheritance. Recent studies have indicated that the regulation and function of nuclear RNAi-mediated heterochromatin are highly complex. Our knowledge of histone modifications and the corresponding histone modifying enzymes involved in the system remains limited. In this study, we show that the heterochromatin mark, H3K23me3, is induced by nuclear RNAi at both exogenous and endogenous targets in *C. elegans*. In addition, dsRNA-induced H3K23me3 can be inherited for four generations. We demonstrate that the histone methyltransferase SET-32, methylates H3K23 *in vitro*. Both *set-32* and the germline nuclear RNAi Argonaute, *hrde-1*, are required for nuclear RNAi-induced H3K23me3 *in vivo*. Our data poise H3K23me3 as an additional chromatin modification in the nuclear RNAi pathway and provides the field with a new target for uncovering the role of heterochromatin in transgenerational epigenetic silencing.

## Introduction

Nuclear RNAi is an evolutionarily conserved pathway in which small RNA mediates transcriptional silencing and heterochromatin formation ^1-4^. Nuclear RNAi plays an important role in genome stability and germline development. It is a highly tractable system for the study of RNA-mediated chromatin regulation and epigenetic inheritance.

*C. elegans* provides a number of unique advantages for the study of nuclear RNAi ^5^. Genetic screens have identified numerous protein factors involved in this pathway. We and others have characterized over 150 genomic loci in *C. elegans* that are de-silenced and/or lose repressive chromatin modifications in nuclear RNAi-deficient mutants, the so-called “endogenous targets” ^6,7^. In addition to these naturally occurring silencing events at the endogenous targets, which include transposons and other “non-self” genes. Nuclear RNAi can also be experimentally triggered at actively transcribed genes by exogenous dsRNA administration ^8,9^, and at germline genes can induce heritable silencing. In the *C. elegans* germline, nuclear RNAi relies on the Argonaute protein, HRDE-1 ^8,10-13^. In current models, HRDE-1 binds secondary siRNAs and recruits nuclear RNAi factors, including chromatin modifying enzymes and remodeling factors, to genomic sites of RNAi ^14^. Germline nuclear RNAi-deficient mutants in *C. elegans* exhibit several phenotypes, including progressive sterility under heat stress (Mrt phenotype) and large-scale de-silencing and chromatin decompaction at the endogenous targets ^5-8,10,12,13,15-17^. We and others have observed the intricacies of this chromatin decompaction (explained below and in the discussion), the consequences are, as yet, unknown.

There are two known nuclear RNAi-induced histone modifications: trimethylation at lysine 27 and lysine 9 of histone H3 (H3K27me3 and H3K9me3), the best studied of which is H3K9me3 ^9,18^. While the enzymes responsible for H3K9me3 have been studied, the function of this histone modification in nuclear RNAi remains unknown. In nuclear RNAi, it was assumed that heterochromatin was required for transcriptional silencing, surprisingly, H3K9me3 is not essential for silencing maintenance if HRDE-1 is present ^7,10,19-21^. By uncovering the histone methyltransferases (HMTs) responsible for each histone modification we can better understand their function in nuclear RNAi. Two HMTs, MET-2 and SET-25, are suggested to function sequentially for H3K9 methylation ^22^. In embryos, MET-2 and SET-25 appear to be the sole contributors of H3K9 methylation ^22,23^. However, in adults, nuclear RNAi-mediated H3K9me3 is dependent on a third HMT, SET-32, in addition to MET-2 and SET-25 ^7,18,24^. While MET-2, SET-25 and SET-32 are all required for the formation of H3K9me3, they are not functionally equivalent. MET-2, but not SET-25, is required for DNA replication stress survival ^20,25^. SET-25 and SET-32, but not MET-2, are required for the silencing of a piRNA-targeted reporter gene ^10^. In addition, while MET-2, SET-25, and SET-32 were all dispensable for silencing maintenance in nuclear RNAi, SET-32 and, to a lesser extent, SET-25 are required for the silencing establishment ^15,21^. Given SET-32’s unique role in nuclear RNAi, we hypothesized that its biochemical activity may differ from MET-2 and SET-25.

The chromatin landscape in *C. elegans* is dynamically regulated during both somatic and germline development ^26,27^. From the embryonic stage to adulthood the two most prominently methylated lysines of histone H3 are H3K27 and H3K23, while H3K9me is proportionally much lower ^26,28^. H3K23me3 has been suggested as a heterochromatin mark in *C. elegans* ^26,28^ and *Tetrahymena* ^29^ and is involved in DNA damage control ^29^. In comparison to the two classical heterochromatin marks, H3K9me and H3K27me, H3K23me is poorly studied. Histone lysine methylation, with the exception of H3K79, is catalyzed by SET-domain containing histone methyltransferases ^30-32^. Although different HMTs share the core catalytic motifs in the SET domain, they can target different lysine residues with high specificity ^32^. The SET-domain containing EZL3 in *Tetrahymena* is required for H3K23me3 *in vivo* ^29^. In *C. elegans*, loss of SET-32 causes reduced levels of H3K23me1 or H3K23me2, however, H3K23me3 was not tested ^21^. No H3K23 HMT has been biochemically validated at the time of this manuscript.

In this paper we enzymatically characterized SET-32 and determined that it is an H3K23 methyltransferase *in vitro*. We show that H3K23me3 can be induced by exogenous dsRNA and is heritable for four generations. H3K23me3 also occurs at the endogenous targets of nuclear RNAi. In addition, H3K23me3 at nuclear RNAi targets is dependent on HRDE-1 and SET-32, and, to a lesser extent, MET-2 and SET-25.

## Results

### SET-32 methylates lysine 23 of histone H3 *in vitro*

To determine the enzymatic activity of SET-32, we performed histone methyltransferase (HMT) assays using recombinant GST-SET-32 and [3H]-labeled S-adenosylmethionine (SAM) (Figure S1). We first tested SET-32’s ability to methylate each of the four core histone proteins, and found that GST-SET-32 methylated free and nucleosomal H3, but not H2A, H2B, or H4 (Figure 1A). There are four conserved catalytic motifs in the SET-domain family proteins, all of which are found in SET-32 (Figure S2) ^32^. The highly conserved tyrosine residue at position 448 in motif IV is predicted to be one of the catalytic residues of SET-32. In order to determine if this residue is required for SET-32’s HMT function, we mutated this tyrosine to phenylalanine (Y448F). The Y448F mutation abolished the HMT activity of GST-SET-32 on H3 (Figure 1B).

**Figure 1:**
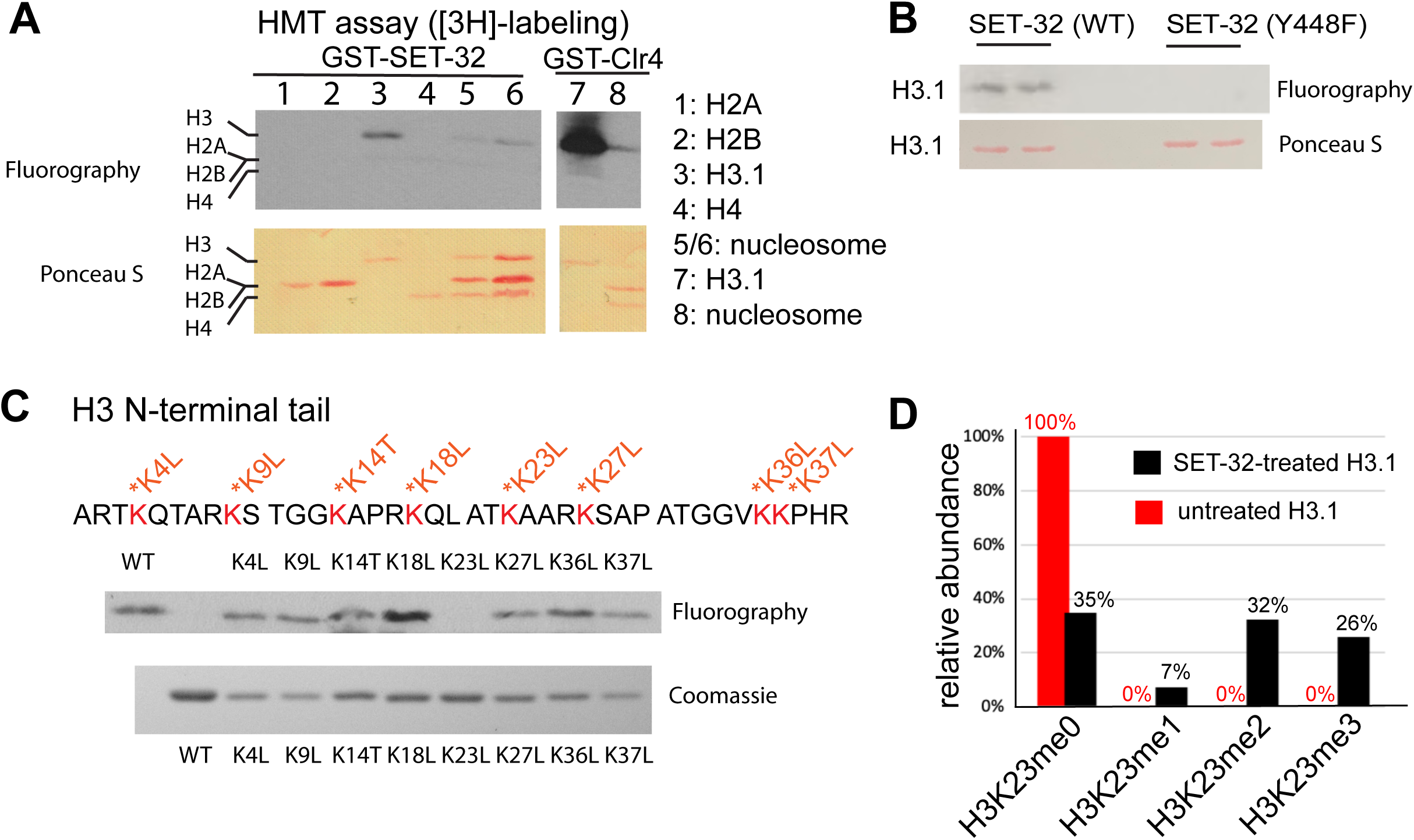
SET-32 methylates H3K23 *in vitro*. **(A)** Detecting the HMT activity of GST-SET-32 by [3H]-labeling and fluorography. Individual core histone proteins and *in vitro* assembled mononucleosome made of 601 DNA^52^ and recombinant *C. elegans* H2A, H2B, and H3.1, and *Xenopus H4*. Xenopus H4 was used because *C. elegans* H4 expression was not successful and there is only one amino acid difference between the two. GST-Clr4 was used as a positive control. **(B)** HMT assay of wild type GST-SET-32 and the GST-SET-32 (Y448F) mutant protein using histone H3.1. **(C)** HMT assay of GST-SET-32 using WT H3.1 and eight lysine mutants of H3.1. **(D)** Mass spectrometry analysis of GST-SET-32-treated H3.1 versus untreated H3.1. The percentage of H3K23-containing fragments with H3K23me0, 1, 2, and 3 is indicated above bars.

The free N-terminal tail domain of H3 contains eight lysine residues available for methylation. To determine whether any of these lysines is required for GST-SET-32’s HMT activity, we generated eight mutant H3 proteins. In each mutant we substituted one of the eight lysines to either leucine or threonine (Figure 1C). GST-SET-32 was able to methylate every mutant H3, except for H3K23L (Figure 1C), suggesting that the lysine 23 is SET-32’s target. We then performed mass spectrometry on GST-SET-32-treated H3, and detected mono, di, and tri-methylation at the K23 position (Figure 1D). The untreated control H3 had no H3K23 methylation. We did not detected methylation at any of the other lysine residues of H3 (data not shown). These results agree with published histone mass spectrometry analysis of set-32 mutant animals ^21^. Our data indicate that SET-32 has H3K23 methyltransferase activity *in vitro*.

### Nuclear RNAi triggers heritable H3K23me3 at germline genes

Since SET-32 is a nuclear RNAi factor, we hypothesized that H3K23me would be induced by nuclear RNAi. Nuclear RNAi can be triggered by both exogenous dsRNA and mediated by siRNAs at the endogenous targets. We first tested whether H3K23me3 could be induced exogenously, as follows. RNAi in *C. elegans* was induced by feeding worms with *E. coli* expressing dsRNA that is homologous to a target gene. We chose a well characterized germline-specific gene, *oma-1*, as the target gene. Wild type animals were fed with *oma-1* dsRNA, control dsRNA (GFP), or no dsRNA for 3-4 generations, and the synchronized young adult animals were then collected for ChIP-seq. All antibodies used in this study were validated using western blot and/or immunofluorescence analysis (Figure S4). In WT animals, *oma-1* dsRNA feeding triggered no significant increase in H3K23me1, a modest increase in H3K23me2, and a dramatic increase in H3K23me3 (Figure 2A). The peak of H3K23me3 corresponds to the trigger region of the *oma-1* dsRNA, and spreads approximately 0.5kb upstream and 1kb downstream of the *oma-1* gene boundaries. The GFP dsRNA and no dsRNA controls did not show any H3K23 methylation at *oma-1*. In order to compare H3K23me3 with a previously described dsRNA-induced histone mark, we performed side-by-side analysis of H3K23me3 and H3K9me3 and observed closely overlapping peak and spread (Figure S5). To verify that H3K23me3 can be induced at other genes, we fed worms with *smg-1* dsRNA and confirmed that H3K23me3 was enriched at the target chromatin in response to *smg-1* dsRNA, with a profile similar to H3K9me3 (Figure 2C).

**Figure 2.**
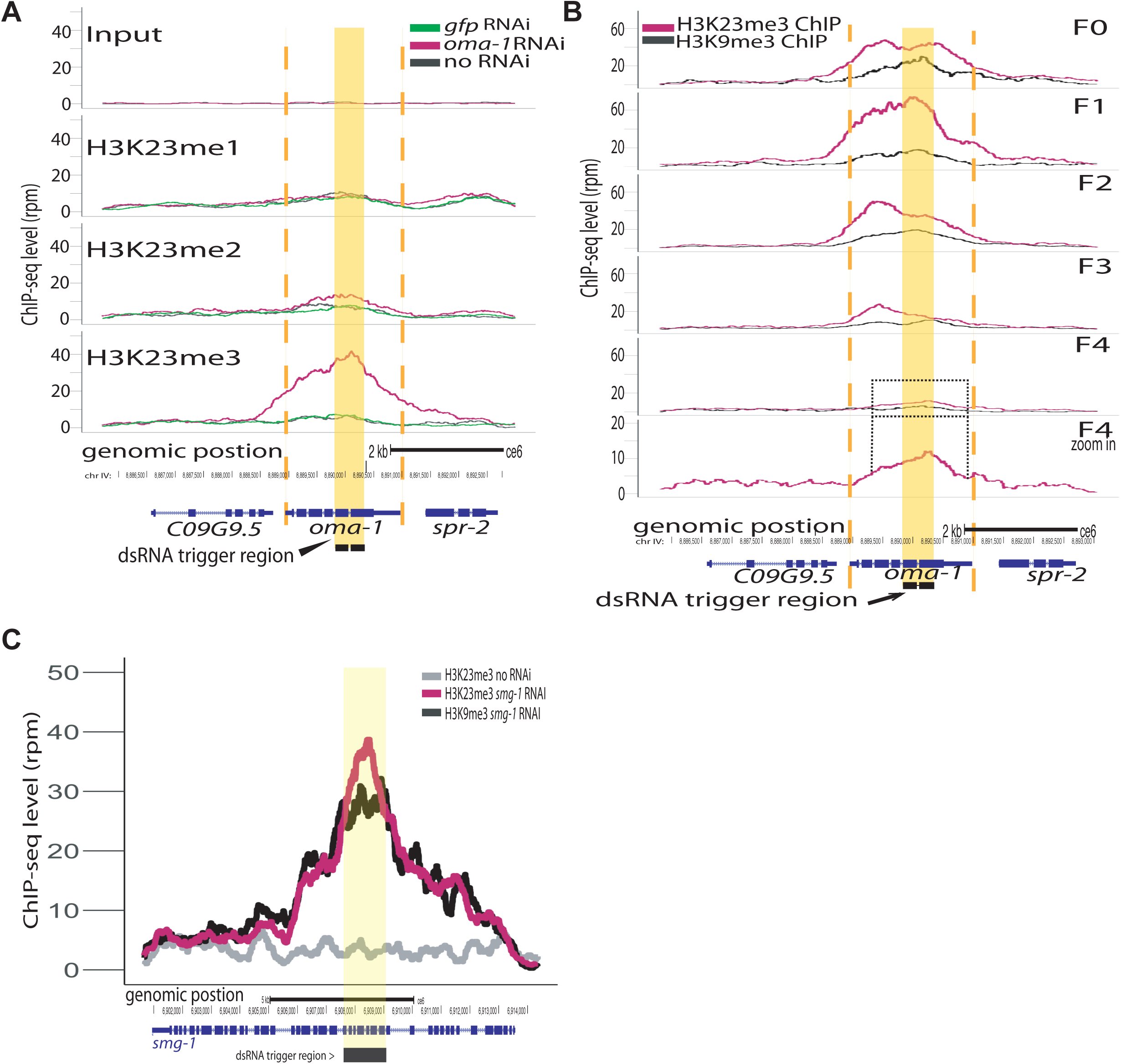
dsRNA triggers heritable H3K23me3 at the RNAi target gene. **(A)** H3K23 methylation levels are plotted as a function of position along the *oma-1* locus. The top panel shows input DNA for ChIP experiments, H3K23me1 (second panel), H3K23me1 (third panel), H3K23me3 (bottom panel), *pink oma-1* dsRNA, *green* GFP dsRNA, *black* no dsRNA feed. **(B)** *oma-1* heritable RNAi assay. H3K23me3 (*pink*) compared with H3K9me3 (*black*), at *oma-1* locus with *oma-1* dsRNA feeding at the F0 generation (top panel) and no dsRNA feeding in subsequent generations, F1-F4. Bottom panel is also F4 but with increased scale to show the continued presence of H3K23me3. Yellow block highlights dsRNA trigger region, orange dashed lines indicate the boundaries of *oma-1*. **(C)** H3K23 and H3K9 methylation levels are plotted as a function of position along the *smg-1* locus after *smg-1* dsRNA feeding (*grey* = no dsRNA feeding). Yellow block highlights dsRNA trigger region. All signals are normalized to sequencing depth.

In order to examine the transgenerational inheritance of dsRNA-induced H3K23me3, we performed a heritable RNAi experiment of *oma-1.* WT animals were fed *oma-1* dsRNA for three generations and subsequently moved to dsRNA-free plates and collected for four generations. As expected, the F0 (dsRNA+) and F1 (dsRNA-) generations exhibited high levels of H3K23me3 and H3K9me3 at the *oma-1* locus (Figure 2B). At F2 (dsRNA-) the H3K23me3 levels remain quite high. By the F3 (dsRNA-), H3K9me3 enrichment fell by two thirds and dropped to the background level at F4 (dsRNA-). The H3K23me3 signal remained robust for up to four generations without dsRNA exposure (Figure 2B). These results indicate that RNAi-mediated H3K23me3 is transgenerationally heritable in *C. elegans*.

### H3K23me3 is a heterochromatic mark in *C. elegans*

In order to further characterize H3K23me3 in *C. elegans*, we conducted ChIP-seq in WT animals and performed whole-genome analysis. The genomic distributions of H3K23me2 and H3K23me3 were highly similar to the genomic distribution of H3K9me3 (Figure 3A & S6). In whole-chromosome coverage plots H3K23me2 and H3K23me3 are enriched where constitutive heterochromatin domains are clustered in the *C. elegans* genome: at the left and right arms of the five autosomes and the left tip of the X chromosome, with the highest peaks at the meiotic paring centers (Figure 3A & Figure S6). H3K23me1 had a relative uniform distribution in the genome (Figure S6). These results were consistent with previous reports that H3K23me2 and H3K23me3 are constitutive heterochromatin marks ^26,28,29^. In order to further assess the correspondence of H3K23me3 and H3K9me3, we plotted both modifications on a scatter plot and observed a high correlation (Pearson coefficient = 0.8) (Figure 3B). Consistent with previous reports, our data shows that H3K23me3 is heterochromatic in distribution and correlates highly with H3K9me3.

**Figure 3.**
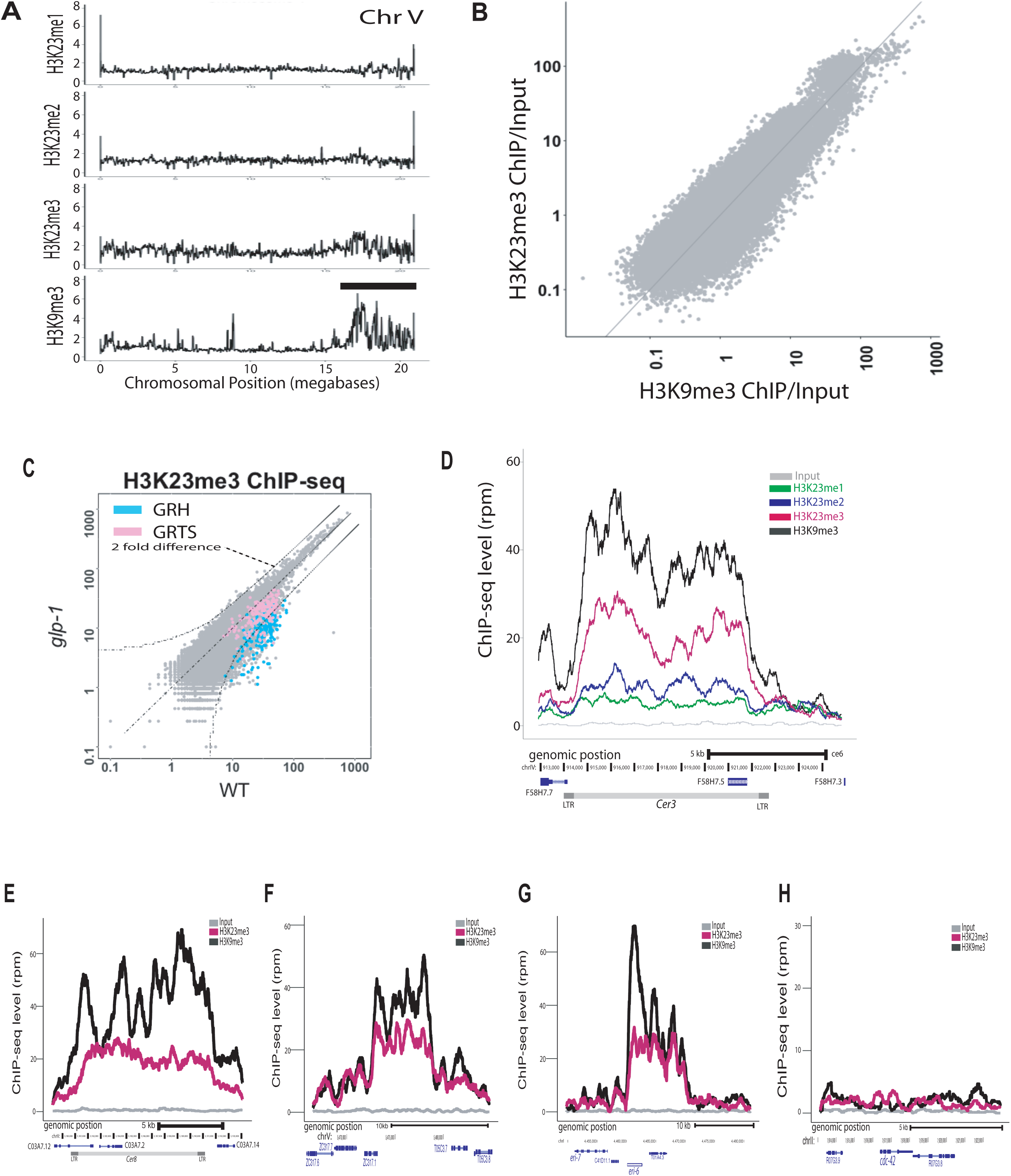
H3K23 methylation profiles at endogenous germline nuclear RNAi targets in WT. **(A)** Relative enrichment (y axis) of H3K23me1, H3K23me2, H3K23me3, H3K9me3 (top to bottom) to input for chromosome V (x axis). Black bar indicates approximate location of meiotic paring center. **(B)** Scatter plot of H3K23me3 ChIP/input (y axis) vs H3K9me3 ChIP/input (x axis) for whole-genome coverage, in which each point represents 1kb of the genome. Averaged values from two replicates were used. **(C)** Scatter plot comparing the H3K23me3 whole-genome profiles (1kb windows) in *glp-1(e2141)* and WT adult animals (25°C). Curved dashed lines indicated two-fold difference (FDR≤0.05). Regions of germline nuclear RNAi-mediated heterochromatin (GRH) are highlighted in pink and regions of germline nuclear RNAi-mediated transcriptional silencing (GRTS) in blue. **(D)** H3K23me1, H3K23me2, H3K23me3, and H3K9me3 enrichment (y axis) at an endogenous germline nuclear RNAi target, *Cer3* LTR retrotransposon (x axis). Grey is the input signal. **(E-H)** H3K9me3 (*black*) and H3K23me3 (pink) coverage plots for three other endogenous targets, **(E)** *Cer8*, **(F)** an exemplary GRH locus on chromosome V:5465000-5485000, **(G)** *eri-6*, and **(H)** a control euchromatin locus, *cdc-42*. All signals are normalized to sequencing depth.

### H3K23me3 is enriched at endogenous targets of nuclear RNAi

We have previously described a set of endogenous targets of germline nuclear RNAi which consist mainly of LTR retrotransposons and other repetitive genomic elements. These HRDE-1-dependent loci are transcriptionally silenced and enriched for H3K9me3. We examined several of these loci for H3K23 methylation using ChIP-seq (Figure 3D-G). At the LTR retrotransposon *Cer3*, H3K23me1 signal was double that of the input, however the signal was still relatively low (Figure 3D). We observed a stronger signal for H3K23me2 (Figure 3D) and a robust signal for H3K23me3 across the endogenous targets, which closely resembles the H3K9me3 signal (Figure 3D-G and data not shown). By contrast, the actively transcribed *cdc-42* locus did not show any enrichment of either H3K9me3 or H3K23me3 (Figure 3H).

As nuclear RNAi is heritable and requires germline factors, we also wished to assess the whether H3K23me3 is a germline-specific histone modification. The *glp-1(e2141)* mutants are defective in germ cell proliferation at the restrictive temperature (25°C). By comparing the *glp-1* mutant and WT worms, we were able to determine that nuclear RNAi-induced H3K23me3 is indeed greatly enriched in the germline (Figure 3C). However, H3K23me3 at both nuclear RNAi targets and globally was still present in the *glp-1* mutant animals, indicating that H3K23me3 occurs in both soma and germline.

By analyzing both exogenous and endogenous targets, we determined that H3K23me3, like H3K9me3, is a nuclear RNAi-induced heterochromatic mark.

### *set-32* and *hrde-1* are required for nuclear RNAi-induced H3K23me3

In order to elucidate the genetic requirements of nuclear RNAi-induced H3K23me3, we performed dsRNA feeding against *oma-1* in three different mutant strains, *hrde-1, set-32 single mutant, and met-2 set-25* double mutant, followed by H3K23me3 ChIP-seq (Figure 4A). As expected, *hrde-1* and *set-32* mutant worms showed greatly reduced H3K23me3 at the *oma-1* locus compared to WT. By contrast, *oma-1* RNAi in the double HMT mutant, *met-2 set-25*, and the WT animals induced similar, high levels of H3K23me3, indicating that SET-32 has a specific role in H3K23 methylation not shared by the other two HMTs (Figure 4A).

**Figure 4.**
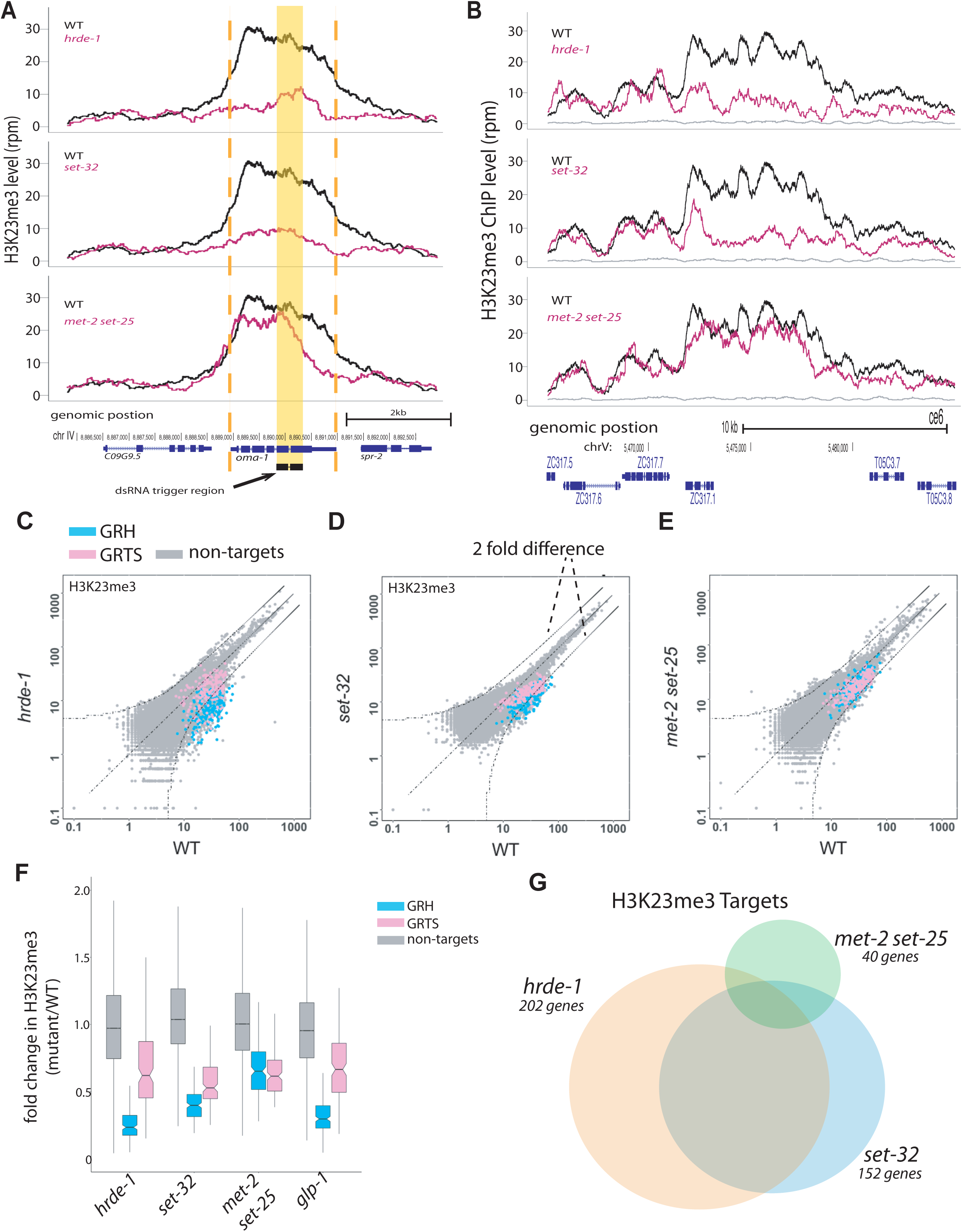
*set-32* and *hrde-1* are required for nuclear RNAi-dependent H3K23me3. **(A-B)** H3K23 methylation levels (y axis) are plotted as a function of genomic position (x axis) in three mutant strains **(A)** for exogenous dsRNA-induced nuclear RNAi at *oma-1* and **(B)** at an endogenous target at chromosome V:5465000-5485000. Top panel: *hrde-1*, middle panel: *set-32*, bottom panel: *met-2 set-25*. The same H3K23me3 signal from WT animals (*black*) was plotted in each panel to compare with the mutant signals (*pink*). Grey: ChIP input from WT. Yellow block highlights the dsRNA trigger region, orange dashed lines indicate the boundaries of *oma-1.* **(C-E)** Scatter plots of H3K23me3 ChIP whole-genome coverage in which each point represents a 1kb window of the genome with GRH and GRST loci highlighted. Mutant coverage is plotted on the y axis and WT coverage is plotted on the x axis. Curved dashed lines indicated two-fold difference (FDR≤0.05). **(F)** A box plot comparing WT and various mutants for GRH and GRTS regions, as well as the rest of the genome. **(G)** Venn diagram of genes enriched with *hrde-1* (orange), *set-32* (blue), and *met-2 set-25* (green)-dependent H3K23me3. Dependence is measured as a 2-fold decrease in H3K23me3 signal compared with WT (p value <0.05) for individual annotated protein-coding genes in two replicas. Fisher’s exact test found the overlap between all pairs are statistically significant; the p values are < 2.2 ×10^−16^ for *hrde-1* vs *set-32* and *set-32* vs *met-2 set-25* and 2.095 ×10^−15^ for *hrde-1* vs *met-2 set-25*.

In order to better understand the contribution of the three HMTs and HRDE-1 to H3K23me3, we took a whole-genome approach (Figure 4). Previous studies have suggested that SET-32 is a nuclear RNAi-specific factor, we therefore expected that the regions of SET-32-dependent H3K23me3 would overlap with HRDE-1 targets. To probe this question, we identified genes with SET-32 or HRDE-1-dependent H3K23me3 profiles. The lists of the top SET-32 and HRDE-1-dependent genes are heavily populated by our previously defined endogenous targets and contain several classes of repeat elements (Table S1 & Figure S8F-H for example coverage plots). We then performed a Venn diagram analysis and found that the majority of SET-32-dependent genes overlapped with HRDE-1-dependent genes (Figure 4G). While there are over 150 SET-32-dependent genes, H3K23me3 is only dependent on MET-2 SET-25 at 40 genes. Our whole-genome scatter plots and box plot showed a strong loss of H3K23me3 in both the *hrde-1* and *set-32* mutant animals and much weaker H3K23me3 reduction in the *met-2 set-25* double mutant animals (Figure 4C-F). As in published works, we found that H3K9me3 has a larger requirement of MET-2 and SET-25 than SET-32 (Figure S7), further supporting the different functions of SET-32 and MET-2/SET-25 ^7,24^.

Interestingly, we did not observe complete abolishment of H3K23me3 in the *set-32* or *hrde-1* mutant animals at either the global level (Figure 4C-D) or in coverage plots (Figure 4B & Figure S8), indicating the existence of additional nuclear RNAi factors and HMTs that contribute to H3K23me3. Although the *met-2 set-25* double mutations had a weaker impact on H3K23me3 than the *hrde-1* and *set-32* single mutation at the global level (Figure 4C-F), the loss of H3K23me3 in *met-2 set-25* mutant varied substantially among different endogenous targets (e.g. comparing Figure 4B and Figure S8A-D). However, when we examined the loss of H3K9me3 in *met-2 set-25* mutant at the same genomic loci we do not see the same variation (Figure S9A-D). These data indicate a complex regulation of heterochromatin marks at the endogenous targets, and suggest the genetic requirements may be contingent on local chromatin environment and other factors.

## Discussion

### *C. elegans* as a model system to study H3K23 methylation

H3K23 methylation was discovered in alfalfa in 1990 ^33^, and subsequently found in yeast, *Tetrahymena, C. elegans*, mouse and human ^21,26,28,29,33-40^. Despite its high degree of evolutionary conservation, H3K23me remains an understudied histone modification. Our knowledge of H3K23me in animals mostly come from *C. elegans*. H3K23 is the second most highly methylated lysine of H3 in *C. elegans* ^29^. The developmental dynamics of H3K23me have been characterized, either alone or in combination with other histone modifications ^26,29^. The whole-genome distributions of H3K23me have been determined in this work and previous studies ^3^. In addition, this study discovered the first biochemically validated H3K23 histone methyltransferase (HMT) and demonstrated that H3K23me3 can be experimentally induced at RNAi target genes. These advances uniquely position *C. elegans* as a powerful system to explore the regulation and function of H3K23 methylation.

### What is the function of H3K23me3 in nuclear RNAi?

We have not yet discovered the functions of H3K23me3 in nuclear RNAi. The co-occurrence of H3K9me3 and H3K23me3 could indicate that they work together to play a larger role in heterochromatin architecture. Both H3K9me3 and H3K23me3 have been shown to bind HP1 or its orthologs *in vitro* ^41^. In addition, both HP1α and H3K23me2 are enriched in the nuclear pore complex in HeLa cells (H3K23me3 was not tested) ^36^.

The involvement of SET-32 in both the maintenance and establishment phases of nuclear RNAi ^7,15,21,24^ suggests that H3K23me3 functions in these phases as well. During the establishment phase of nuclear RNAi, the host organism encounters a foreign genetic element for the first time and must repress its active transcription. In the maintenance phase, a stable silencing state is passed on from parent to progeny.

#### Maintenance

Although SET-32 is dispensable for silencing maintenance when in the presence of wild type HRDE-1, *set-32 hrde-1* double mutants show enhanced de-silencing when compared to *hrde-1* single mutants ^15^. These results suggest a conditional requirement of H3K23me3 for silencing maintenance, possibly to function as a failsafe or backup silencing mechanism.

#### Establishment

Recent studies have shown that SET-32 is required for silencing establishment at both exogenous dsRNA targets and endogenous targets ^15,21^. Unlike in the maintenance phase, the requirement of SET-32 in silencing establishment is not conditional on the lack of HRDE-1, suggesting that H3K23me3 plays a more prominent role in the establishment phase than the maintenance phase.

H3K23me3 is not limited to the endogenous nuclear RNAi targets. Similar to H3K9me3, H3K23me3 is broadly enriched in *C. elegans* heterochromatin. Our studies showed that H3K23me3 in the non-nuclear RNAi regions is independent of SET-32. Future studies are needed to identify additional H3K23 HMTs and the broader function of H3K23me.

### Nuclear RNAi mark induces multiple heterochromatin marks

Previous studies identified H3K9me3 and H3K27me3 as nuclear RNAi-mediated histone modifications ^9,18^. This study adds H3K23me3 to the list. Based on these results, we propose that the revised model should include the following features. (1) Different marks at nuclear RNAi targets are deposited by different HMTs, MET-2 and SET-25 for H3K9me3, MES-2 for H3K27me3, and SET-32 and other HMT(s) for H3K23me3. (2) That SET-32 is required for H3K9me3 suggests that H3K23me3 promotes H3K9me3. (3) At some of the endogenous nuclear RNAi targets, H3K9me3 and H3K23me3 appear to be a mutually dependent. (4) H3K27me3 is largely independent of H3K9me3 and H3K23me3 ^7^. Future studies are needed to further delineate the relationship of these histone marks and determine the mechanisms by which the HMTs are recruited to the target chromatin.

## Acknowledgements

We thank Danesh Moazed, Ruth Steward, James Millonig, Michael Verzi, and Vincenzo Pirrotta for help and suggestions. Research reported in this publication was supported by the Rutgers Busch Biomedical Grant to SSG, the National Institute of General Medical Science of the NIH, United States, under award R01GM111752 to SSG and the New Jersey Commission on Cancer Research under award DCHS19PPC030 to LSO. The content is solely the responsibility of the authors and does not necessarily represent the official views of the NIH.

## Author Contributions

Conceptualization, LSO, JZN and SGG; Methodology, LSO, JZN, SS and SGG; Investigation, LSO, JZN, CZ, RA, DK, AZ, SS, and SGG; Writing, LSO and SGG.

## Materials and Methods

### Plasmid construction and recombinant protein purification for GST-fusion proteins

The pGEX-6P-1-GST-SET-32-WT (pSG361) and pGEX-6P-1-GST-SET-25-WT (pSG355) were generated by inserting *set-32* and *set-25* cDNA fragments into the plasmid pGEX-6P-1 using the BamHI and NotI sites. The *set-32* cDNA fragment was amplified by RT-PCR using *C. elegans* (N2) mRNA. The codon optimized *set-25* cDNA fragment was purchased from IDT as gBlock DNA. To create the pGEX-6P-1-GST-SET-32-Y448F plasmid (pSG434), the AccI-NotI fragment of the pGEX-6P-1-GST-SET-32-WT was replaced with gBlock DNA from IDT carrying the Y448F mutation. Plasmids sequences were confirmed by Sanger sequencing. pGEX-6P-1-GST-Clr4 was a gift from the Danesh Moazed ^42^.

The procedure for protein expression and purification was adapted from ^42^. Briefly, *E. coli* BL21-Gold (DE3) that was transformed with the expression plasmid was cultured in 2xYT at 37°C until OD600 reached 0.6-0.8, followed by incubating on ice for 30 min, and then 30 mins of 18°C in shaker before IPTG induction. Recombinant protein expression was induced by 0.2 mM IPTG and continued overnight in the 18°C shaker. All samples and reagents were placed on ice or in the cold room during protein purification. Cells were collected, resuspended in a lysis buffer (150 mM NaCl, 20 mM sodium phosphate pH=7.4, 1% Triton X-100, 1 mM DTT, and 1 mM PMSF), and lysed by Bioruptor using the high output and five 8-min sessions with 30-sec on/30-sec off cycle. A large fraction of the GST-SET-32 and GST-SET-25 were lost as inclusion bodies. After a clear spin, the soluble GST-tagged protein was pulled down by rotating the sample with glutathione sepharose beads for 1 hour. The beads were washed in a buffer containing 150 mM NaCl, 20 mM sodium phosphate pH=7.4, 1 mM DTT, and 1 mM PMSF three times. Protein was eluted using a buffer containing 50 mM Tris-HCl pH=8.0, 15 mM glutathione, 10% glycerol, 1 mM PMSF, and 1 mM DTT, dialyzed against the storage buffer (50 mM Tris-HCl pH=8.0, 10% glycerol, 1 mM PMSF, and 1 mM DTT), and concentrated using the Vivaspin columns (MWCO 30KDa, GE healthcare). The remaining soluble GST-SET-32 aggregates to form >600 KDa complex as measured by size exclusion chromatography analysis (data not shown).

### Plasmid construction and recombinant protein purification for histone H3 proteins

Plasmid pet28a_human_H3.1 (a gift from Joe Landry, Addgene plasmid # 42631) was used to construct the mutant H3 expression plasmids used in this study. The H3K4L, H3K9L, H3K14T, and H3K18L mutations were introduced by replacing the NcoI-MscI fragment in pet28a_human_H3.1 with NcoI-MscI fragments containing the corresponding mutations. Similarly, the MscI-AgeI fragments were used to make the H3K23L and H3K27L mutations and the AgeI-SalI fragments were used for H3K36L and H3K37L mutations. The single-stranded oligoes or oligo duplex pairs were used to make the mutation-containing fragments. Plasmids pet28a_elegans_H2A (*his-12*, pSG395), pet28a_elegans_H2B (*his-11*, pSG396), pet28a_elegans_H3.1 (*his-9*, pSG397), and pet28a_elegans_H4 (*his-31*, pSG398) were constructed by inserting the cDNA fragments amplified using *C. elegans* mRNA by RT-PCR into the NcoI and NotI sites. The *C. elegans* H2A, H2B, H3.1, and H4 cDNA fragments were amplified by using mRNA isolated from wild type animals. Plasmids sequences were confirmed by Sanger sequencing.

Histone purification was performed using the protocol as previously described in ^43^. Briefly, *E. coli* BL21-Gold (DE3) that was transformed with the expression plasmid was cultured in 2xYT medium at 37°C until OD600 reached 0.6-0.8. After 1 mM IPTG was added, cells were cultured for 4 hours at 37°C. 2 grams of cells were resuspended with 1 ml 10x SA buffer (400 mM sodium acetate, pH 5.2, 10 mM EDTA, 100 mM lysine), 0.5 ml 4M NaCl, 100 μl 100 mM PMSF, 100 μl HALT protease inhibitor, 3.5 μl 2-mercaptoethanol, 3.6 g urea, and diH2O to a final volume of 10 ml. Cells were lysed by Bioruptor using the high output and five 8-min sessions with 30-sec on/30-sec off cycle. After 20 min of spin at 41,000xg, the supernatant was filtered with 0.45 μM syringe filter and passed through the HiTrap Q 5 ml column on a FPLC machine. The flow-through was loaded onto HiTrap SP 5ml column. The elution was done by 25 ml 0-19% buffer B and 50 ml 19-50% buffer B. Buffer A contains 40 mM sodium acetate, pH 5.2, 1 mM EDTA, 10 mM lysine, 200 mM NaCl, 6 M urea, 1 mM PMSF, 5 mM 2-mercaptoethanol. Buffer B has 1 M NaCl and is otherwise the same as buffer A. Histone containing fractions were determined by SDS-PAGE/coomassie analysis. The expression of *C. elegans* H2A, H2B, and H3 was successfully but H4 was not (Figure S3). Xenopus H4 (Histone Source), which is identical to *C. elegans* H4 except at position 74 (threonine in Xenopus and cysteine for *C. elegans*), was used for nucleosome assembly.

### Nucleosome assembly

Nucleosome was assembled as previously described ^43-45^. Briefly, *C. elegans* H2A (12 μM), H2B (12 μM), H3 (10 μM), and Xenopus H4 (10 μM) (Histone Source) proteins were mixed in a 250 μl final volume containing 7 M guanidinium HCl, 20 mM Tris-HCl, pH 7.5, 10 mM DTT. Guanidinium HCl was added directly to the mixture, which was then rotated at room temperature for 30-60 min to dissolve guanidinium and spun to clear the mixture. The supernatant was dialyzed against 1L refolding buffer (2M NaCl, 10 mM Tris-HCl, pH 7.5, 1 mM EDTA, 5 mM 2-mercaptoethanol) three times (2 hours overnight, and 2 hours in the cold room). After a clear spin, the sample was fractionated by size exclusion chromatography (Superdex 200 10/300 GL) in the refolding buffer. The histone octamer fractions were identified by SDS-PAGE/coomassie analysis (Figure S3). 12 μg 601 DNA (Histone Source) in 2 M NaCl was mixed with 12 μg histone octamer in the refolding buffer and then dialyzed against a series of buffers with 10 mM Tris-HCl, pH 7.5, 1 mM EDTA, 1 mM DTT, and reducing amounts of NaCl (1 M, 0.8 M, 0.6 M, and 0.05 M), 2 hours for each dialysis. The reconstituted nucleosome was heated at 55°C for 20 min and then cooled to room temperature for 10 min for most thermal stable nucleosome positioning. The nucleosome was concentrated and used for the HMT assay.

### HMT assay

The HMT assay was performed as described in ^42^. Briefly, the [3H]-based HMT assay was carried out in a 10 µl reaction mix containing 2 µM histone or nucleosome, 5.6 µM [3H]-S-adenosyl methionine (SAM), 0.3-1 µM enzyme, and 1x HMT buffer (50 mM Tris-HCl, pH 8.0, 20 mM KCl, 10 mM MgCl_2_, 0.02% Triton X-100, 1 mM DTT, 5% glycerol, and 1 mM PMSF). The reaction mix was incubated at 20°C for 2 hours, and then loaded onto 17% SDS-PAGE. After electrophoresis, proteins were transferred to PVDF membrane, which was then soaked with autoradiography enhancer (EN3HANCE™, PerkinElmer) and then air dried. Fluorography signal was detected by X-ray film. For the histone mass spectrometry analysis, 150 µl reaction mix containing approximately 0.3 µM GST-SET-32, 2.5 µM H3, and 213 µM SAM and 1 x HMT buffer without Triton X-100 was used.

### Mass spectrometry

Histone peptides were obtained as previously described ^46^. Briefly, histone pellets were resuspended in 20 μL of 50 mM NH_4_HCO_3_ (pH 8.0) plus 5 µl of acetonitrile. Derivatization was performed by adding 5 µl of propionic anhydride rapidly followed by 16 µl of ammonium hydroxide and incubated for 20 min at room temperature. The reaction was performed twice to ensure complete derivatization of unmodified and monomethylated lysine residues. Samples were then dried, resuspended in 20μL of 50 mM NH_4_HCO_3_ and digested with trypsin (Promega) (enzyme:sample ratio= 1:20, 2 hours, room temperature). The derivatization reaction was then performed again twice to derivatize peptide N-termini. Samples were then desalted by using in-house packed stage-tips and dried using a SpeedVac centrifuge.

Dried samples were resuspended in 0.1% trifluoroacetic acid (TFA) and injected onto a 75 µm ID x 25 cm Reprosil-Pur C18-AQ (3 µm; Dr. Maisch GmbH, Germany) nano-column packed in-house using a Dionex RSLC nanoHPLC (Thermo Scientific, San Jose, CA, USA). The nanoLC pumped a flow-rate of 300 nL/min with a programmed gradient from 5% to 28% solvent B (A = 0.1% formic acid; B = 80% acetonitrile, 0.1% formic acid) over 45 minutes, followed by a gradient from 28% to 80% solvent B in 5 minutes and 10 min isocratic at 80% B. The instrument was coupled online with a Q-Exactive HF (Thermo Scientific, Bremen, Germany) mass spectrometer acquiring data in a data-independent acquisition (DIA) mode as previously optimized ^47^. Briefly, DIA consisted on a full scan MS (*m/z* 300−1100) followed by 16 MS/MS with windows of 50 *m/z* using HCD fragmentation and detected all in the orbitrap analyzer.

DIA data were searched using EpiProfile 2.0 and validated manually ^48^. The histone H3 peptide KQLATKAAR (aa 18-26) was considered in all possible modified forms (unmodified, me1/2/3). The relative abundance of each form was calculated using the total area under the extracted ion chromatograms of all peptides in all the (un)modified forms and considered that as 100%. To confirm the position of the methylation, we extracted the chromatographic profile of the MS/MS fragment ions and verified that no unique fragment ions belonging to the K18me1/2/3 possible peptide isoforms had detectable intensity.

### *C. elegans* Strains

*C. elegans* strain N2 was used as the standard WT strain. Alleles used in this study were LG I: *set-32*(red11), LG III: *hrde-1*(tm1200), *glp-1*(e2141), *set-25*(n5021), *set-32*(ok1457), *met-2*(n4256) *set-25* (ok5021). *C. elegans* were culture on NMG agar plates that was as previously described ^49^ in a temperature-controlled incubator.

### Preparation of worm grinds

Synchronized young adult worms were first washed off the plates with M9 buffer. *E. coli* OP50 bacteria washed off together with the worms were separated and removed by loading the worms to 10% sucrose cushion and centrifuging for 1 minute at 600g in a clinical centrifuge. Worms were then pulverized by grinding in liquid nitrogen with a pre-chilled mortar and pestle and were stored at −80°C.

### dsRNA feeding

Worms were grown on NGM plates with the following food sources: OP50 *E. coli* (no RNAi), oma-1 *E. coli* (plasmid SG42), GFP *E. coli* (plasmid SG221), smg-1 *E. coli* (plasmid SG27). Worms were synchronized by bleaching, subsequent starvation and released at the L1 stage onto described plates. Each assay in this study used grinds of ∼5000 worms for each condition. For RNAi experiments, worms were grown for three to four generations on RNAi culture before grinding. For heritable RNAi experiments we used the protocol described in^9^. Briefly, worms were raised on oma-1 RNAi plates for three generations before synchronization and release onto OP50 *E. coli* plates without dsRNA feed. Worms were collected at P0, F1, F2, F3, F4 generations for grinding.

### ChIP-seq library construction

Worm grinds from approximately 5000 worms were used for each chromatin immunoprecipitation experiment according to the procedure described in ^5^. Anti-H3K9me3 (ab8898, Abcam) and anti-H3K23me3 (61500, Active Motif) antibodies were used for the H3K9me3 and H3K23me3 ChIP, respectively. Each ChIP experiments usually yielded 5–10 ng DNA. The entire ChIP DNA or 10 ng DNA in the case of ChIP input was used to make DNA library with the KAPA Hyper Prep Kit (KAPA Biosystems) according to the manufacturer’s instruction. For each sample in a given assay worm grinds were thawed and subsequently crosslinked and sonicated to produce fragments between 200-500bp according to protocol described in ^6^. Samples were then used for ChIP (H3K23me1,2,3, H3K9me3, H3K27me3) or stored for library prep as input DNA. IP was performed with the following antibodies: anti-H3K9me3 (ab8898, Abcam), anti-H3K23me1 (39388, Active Motif), anti-H3K23me2 (39654, Active Motif), anti-H3K23me3 (61500, Active Motif), and anti-H3K27me3 (39535, Active Motif). For each antibody ∼0.5-1.5% of input DNA was pulled down, with DNA yields between ∼5-25 ng. 5 ng or less of DNA was used for library preparation using KAPA Hyper Prep Kit (KAPA Biosystems) according to the manufacturer’s instruction. PCR was performed on library DNA for 12-17 cycles after which all libraries were pooled according to Illumina HiSeq specifications. Sequencing was sent to Illumina and carried out according to the following specifications: 50-nt single-end run, dedicated index sequencing. Dedicated 6-mer indexes were used to demultiplex the libraries of different samples. All libraries used in this study are listed and described in Supplementary Table S2. De-multiplexed raw sequencing data in fastq format for each library is available at NCBI (GEO accession number:).

### Antibody Validation

#### Immunofluorescence staining

Adult worms were washed in 1X PBS twice and paralyzed in 0.1 mM levamisole in 1X PBS. Paralyzed worms were transferred to a cavity slide and gonads were dissected using two 25-gauge syringe needles. The dissected gonads were first fixed in 100% methanol in 1X PBS at - 20ºC for 1 minute, and then were fixed in 2% paraformaldehyde in 1X PBS at room temperature for 5 minutes. After fixation, gonads were blocked in blocking buffer (1 mg/ml BSA, 10% Normal Goat Serum, 0.1% Tween 20, 1X PBS) for 45 minutes at room temperature. For primary antibody staining, gonads were incubated in Rabbit-anti-H3K23me3 (1:150, Active Motif, 61499) in blocking buffer at 4ºC overnight. For antibody competition, primary antibody was pre-absorbed with 25ng/µL corresponding histone proteins or histone peptide at room temperature for 1 hour, and was centrifuged at 14,000rpm at 4ºC for 10mins to remove immune complexes. The histone proteins and peptide used for antibody competition were histone H3K23me3 (Active motif, 31264), histone H3K9me3 (Active motif, 31601), unmodified histone H3 (Abcam ab2903), H3K27me3 histone peptide (Abcam ab1782). For secondary antibody staining, gonads were washed three times for 5 minutes each wash in 1X PBS/0.1% Tween 20, and then were incubated in Donkey Anti-Rabbit-Alexa Fluor^®^ 488 (1:300, Jackson ImmunoResearch Laboratories, 711-545-152) in blocking buffer at room temperature for 2 hours. The gonads were then washed three times for 5 minutes each wash in 1X PBS with 0.1% Tween 20. 100 ng/ml DAPI was added to the last wash to stain chromosomes. Gonads were mounted in Slowfade (Invitrogen) onto a freshly made 2% agarose pad for imaging. Gonads were imaged using a Zeiss Axio Imager M2 system. Images were processed with Fiji (ImageJ)^50^.

#### Western blot

Worm grinds were lysed in 2X Laemmli buffer with 1X HALT protease and phosphatase inhibitor (Thermo Fisher Scientific) by boiling at 95°C for 5 minutes. Worm lysate (25 µg /lane) were loaded and separated on Bio-Rad TGX Any KD gel and transferred to nitrocellulose membrane. Primary antibodies used are polyclonal Rabbit-anti-H3K23me1 (1:1000, Active Motif 30508001), polyclonal Rabbit-anti-H3K23me2 (1:1000, Active Motif 28209001), polyclonal Rabbit-anti-H3K23me3 (1:1000, Active Motif 31913001), polyclonal Rabbit-anti-H3K9me3 (1:1000, Abcam ab8898), monoclonal Mouse-anti-tubulin (1:250, DSHB AA4.3). Secondary antibodies used are Cy5-conjugated donkey anti-Rabbit IgG secondary antibody (1:1000, Jackson ImmunoResearch Laboratories 711-175-152) and Cy5-conjugated donkey anti-Mouse IgG secondary antibody (1:1000, Jackson ImmunoResearch Laboratories, 715-175-150). Fluorometric detection and measurement was preformed using GE/Amersham Typhoon RGB scanner and ImageQuant software (GE Healthcare).

### High-throughput sequencing

Pooled libraries were sequenced on an Illumina HiSeq 2500 platform (rapid run mode, 50-nt single-end run, and index sequencing). De-multiplexed raw data in fastq format were provided by the sequencing service facility. Library information is listed in Supplemental Table S2. High-throughput sequencing data generated for this study have been deposited in NCBI Gene Expression Omnibus (accession number GSE141347).

### Data analysis

Fastq files of each library were aligned directly to *C. elegans* genome (WS190 version) using the Bowtie alignment program (version 1.2.2), only perfect alignments were reported and used ^51^. When a read aligned to multiple loci the alignment was counted as 1/(number alignments). For all data analysis normalization was based on the sequencing depth of each library (total number of reads aligned). For some figures, normalization was additionally based on the respective input library (no antibody), where this is done it is stated in the figure legend. For individual loci coverage, each read was extended by 500 bp from the sequenced end. Whole chromosome coverage analysis was done based on 1kb windows for the entire genome. All coverage plot figures were created using custom python scripts and custom R scripts in ggplot2.

### Venn Diagram Analysis

We used venn.js (https://github.com/benfred/venn.js/), a library in D3.js. To layout each Venn diagram proportional to the input sizes, we defined the sets, and specified the size of each individual set as well as the size of all set intersections. The sizes of all sets and intersections were found in custom R scripts. Each set was defined using two replicate libraries.

## Supplementary figure legend

**Figure S1.**
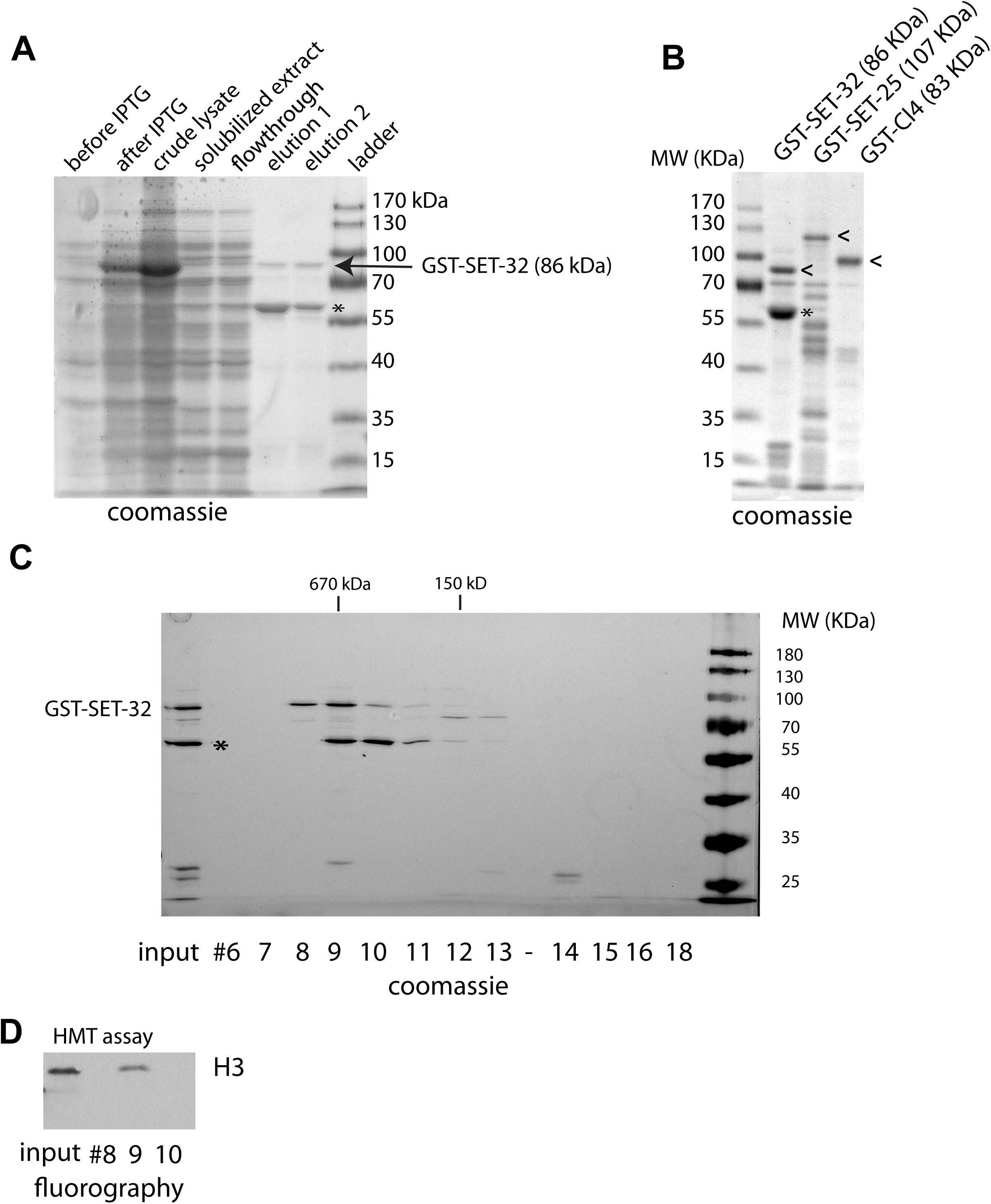
Recombinant GST-fusion protein purification. **(A)** SDS-PAGE/coomassie analysis of GST-SET-32 expression and purification. The strong reduction of GST-SET-32 after clear spin (compare the crude lysate and solubilized extract) indicates that most of the GST-SET-32 was expressed as inclusion body. **(B)** SDS-PAGE/coomassie of the GST-SET-32, GST-SET-25, and SET-Clr4 purification products. The full-length GST fusion proteins are indicated by <. **(C)** SDS-PAGE/silver stain analysis of size exclusion chromatography fractions (Superdex 200 10/300 GL column from GE, 1 ml fractions) of the GST-SET-32 purification product. The main co-purified 60KDa protein (indicated by *) and the GST-SET-32-containing fractions largely overlap. Removing the GST-tag by HRV 3C protease did not change the overlapping of SET-32 and the 60KDa protein in size exclusion chromatography (data not shown). **(D)** Fluorography of HMT assays ([3H]-labeling of H3) using fractions 8, 9, and 10, as well as the input, of the size exclusion chromatography of GST-SET-32. Note that the peak GST-SET-32 fraction (#9) has the HMT activity.

**Figure S2.**
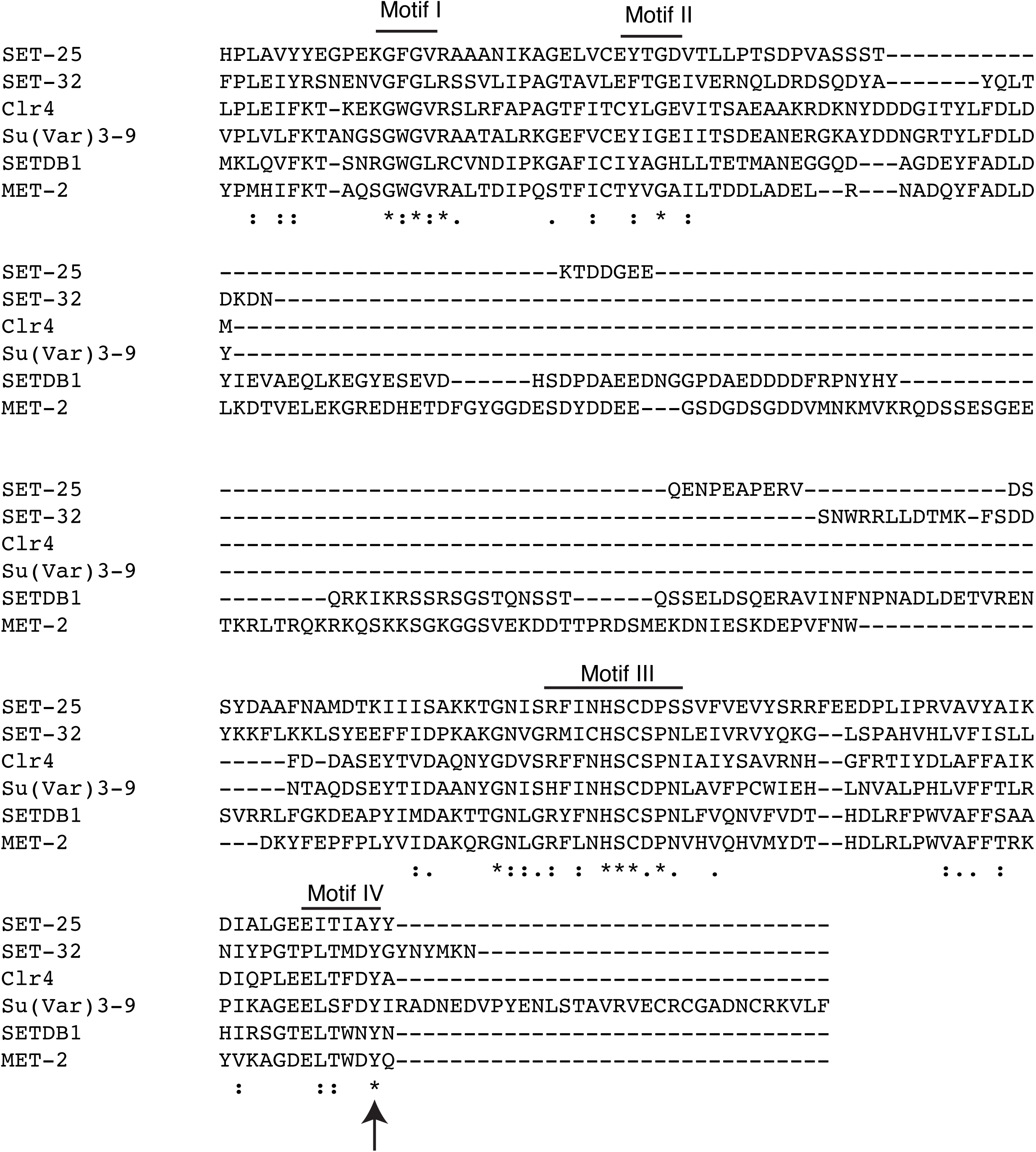
Alignment of SET domains of histone methyltransferases. The four conserved motifs and Y448 (in SET-32) position are indicated. Arrow indicates Y448F residue.

**Figure S3.**
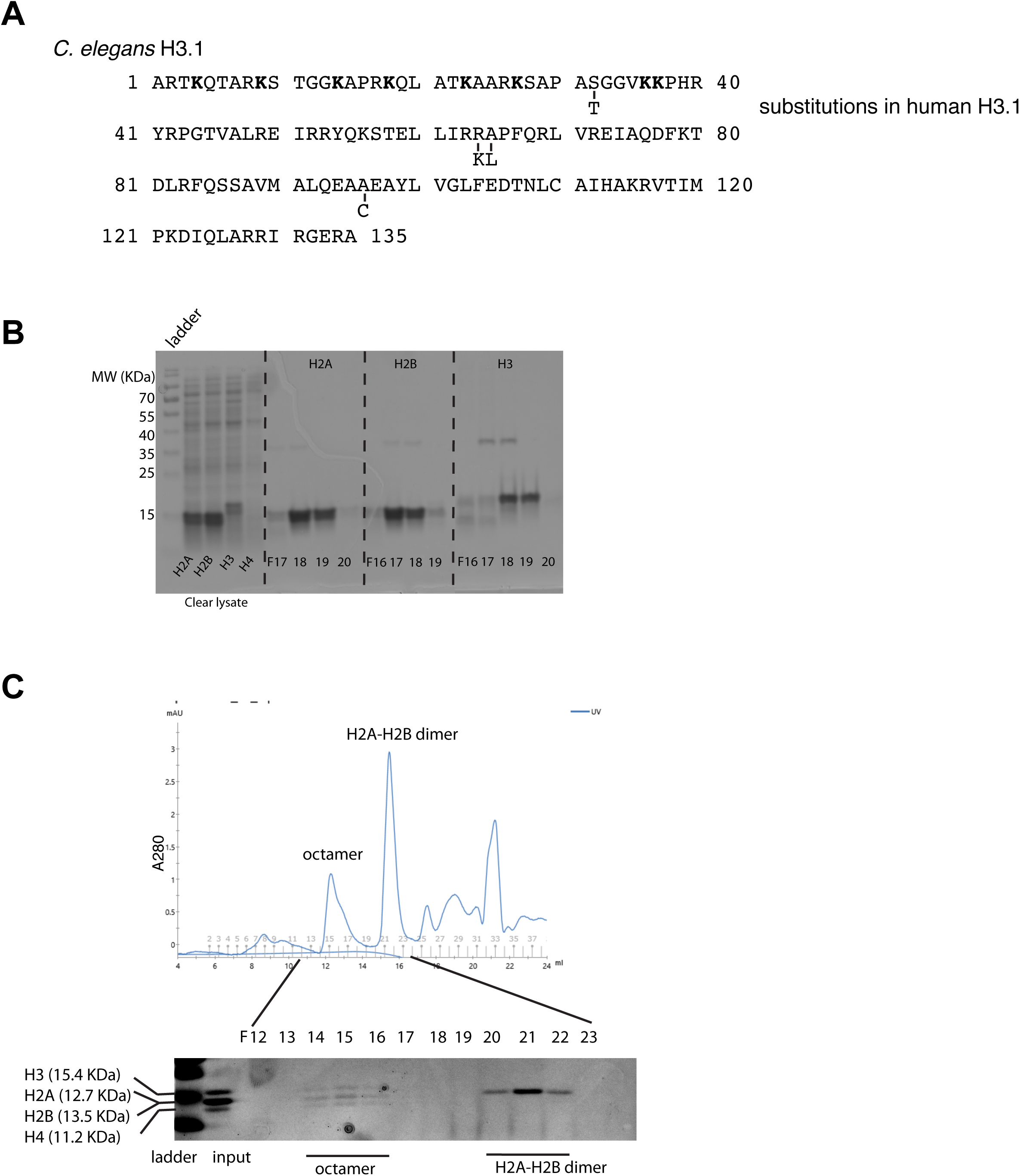
Recombinant histone purification and histone octamer assembly. **(A)** *C. elegans* H3.1 protein sequence, lysine residues within the first 40 residues in bold. The differences with human H3.1 sequence are indicated. **(B)** SDS-PAGE/coomassie analysis of *C. elegans* histone expression and purification. Lysate before HiTrap SF fractionation are shown to indicate expression of H2A, H2B, and H3, but not H4 in *E. coli*. Peak fractions of HiTrap SF chromatography are shown for H2A, H2B, and H3. **(C)** Size exclusion fractionation of histone octamer assembly. *C. elegans* H2A, H2B, H3 and *Xenopus* H4 were used. *Xenopus* H4 (Histone Source) is identical to *C. elegans* H4 except at position 74 (threonine in Xenopus and cysteine for *C. elegans*). (*C. elegans* and *Xenopus* H2Bs have the identical protein sequence. The percentages of identity between the two species are 81.7% and 95.6% for H2A and H3, respectively.)

**Figure S4.**
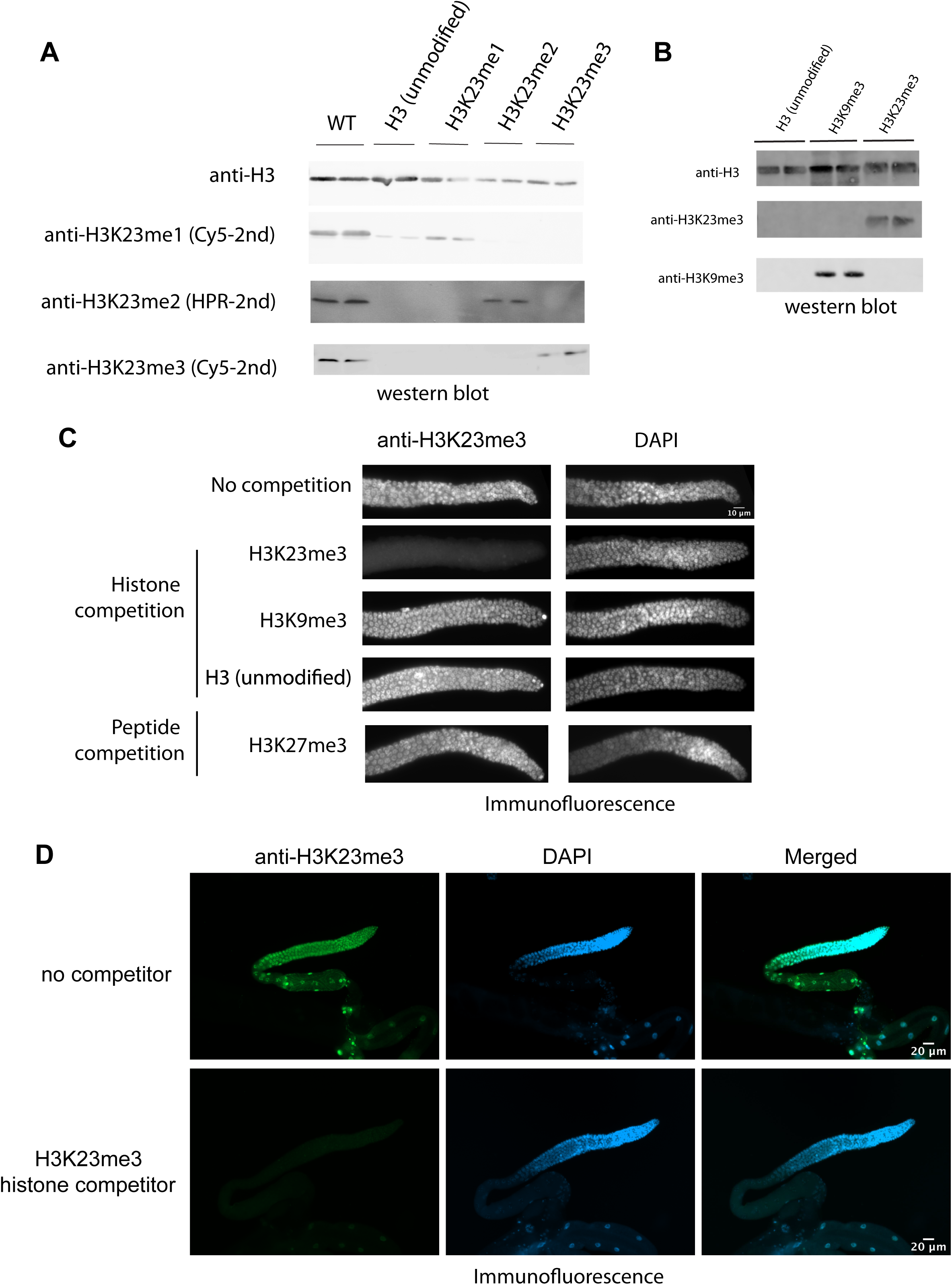
Antibody validation. **(A)** Western blotting against *C. elegans* crude lysate, recombinant H3 with or without H3K23 methylation, as indicated at the top. Antibodies are indicated on the left. **(B)** Western blotting against recombinant H3, H3K9me3, and H3K23me3. **(C)** Immunofluorescence analysis: Anti-H3K23me3 and DAPI staining of dissected gonads from adult *C. elegans*. Scale bar: 10 μm. The antibody was pre-incubated with either H3, H3K23me3, H3K9me3 histone proteins or H3K27me3 peptide, as indicated on the left. Anti-H3 and anti-H3K9me3 antibodies were purchased from Abcam; anti-H3K23me1/2/3 from Active Motif. **(D)** Same assay as figure (C) but with whole gonad shown, anti-H3K23me3 and DAPI staining of whole gonads from adult *C. elegans*. Scale bar: 20 μm. The antibody was pre-incubated with H3K23me3 histone protein in lower panel.

**Figure S5.**
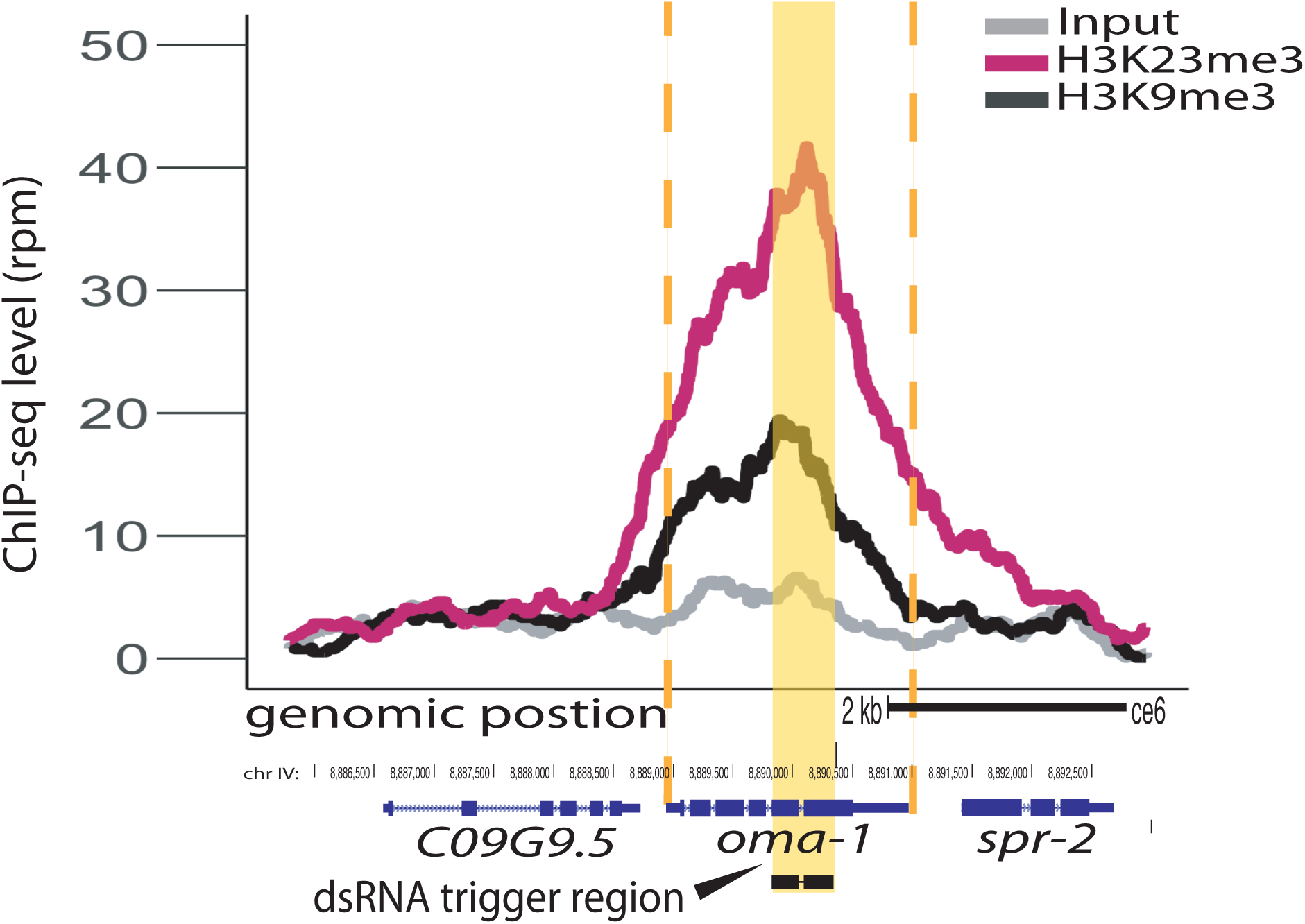
Comparison of dsRNA mediated H3K9me3 and H3K23me3. H3K23me3 (*pink*) compared with H3K9me3 (*black*), at the *oma-1* locus, *grey* no dsRNA feeding. All signals normalized to sequencing depth. Yellow block highlights dsRNA trigger region, orange dashed lines indicate the boundaries of *oma-1*.

**Figure S6.**
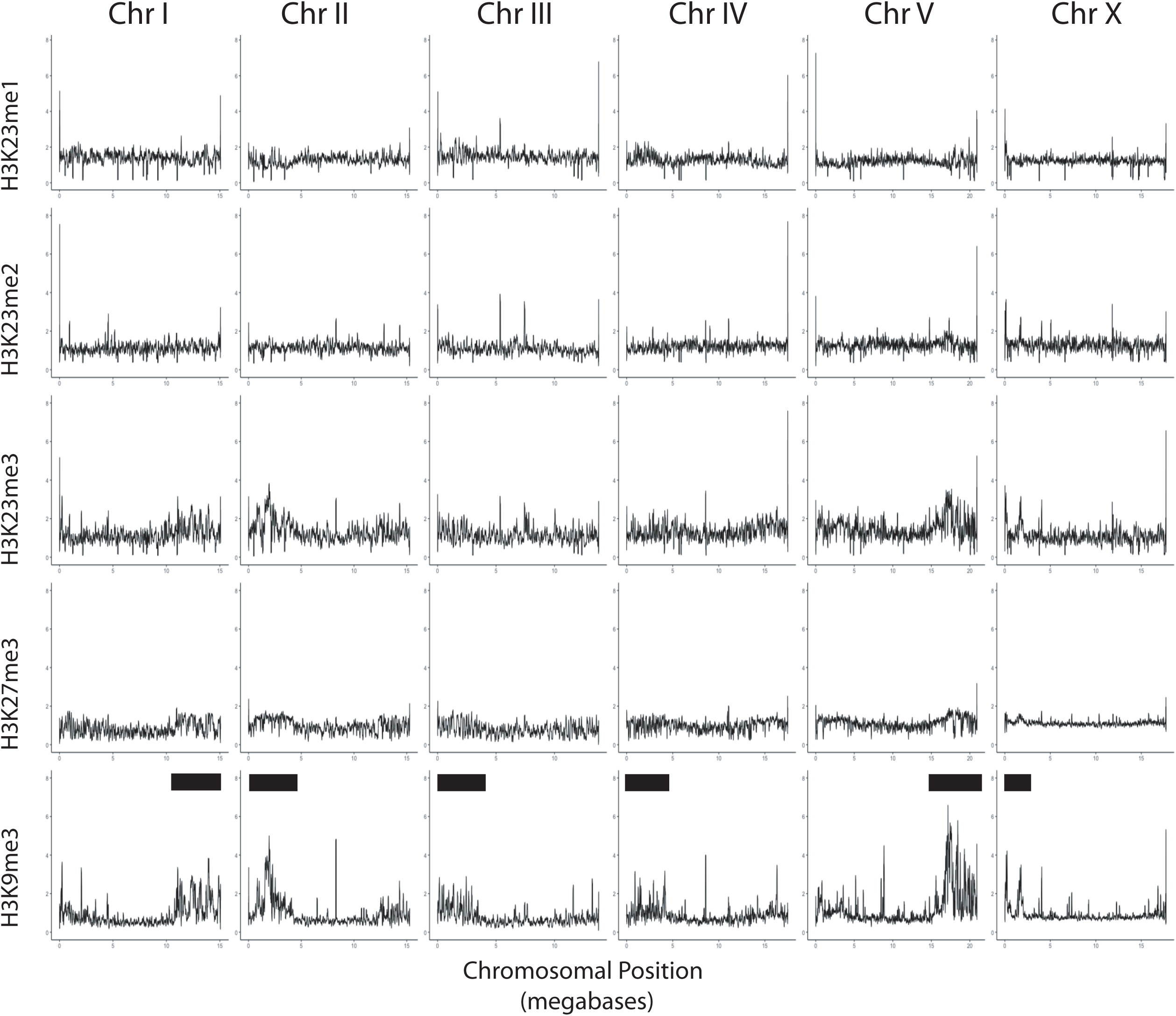
Whole Chromosome Comparison of H3K9me/H3K27me/H3K23me. Relative enrichment of H3K23me1, H3K23me2, H3K23me3, H3K27me3, H3K9me3, (top to bottom) to input (y axis) for each chromosome (x axis). Black bar indicates approximate location of meiotic paring centers. Signal is normalized to sequencing depth.

**Figure S7.**
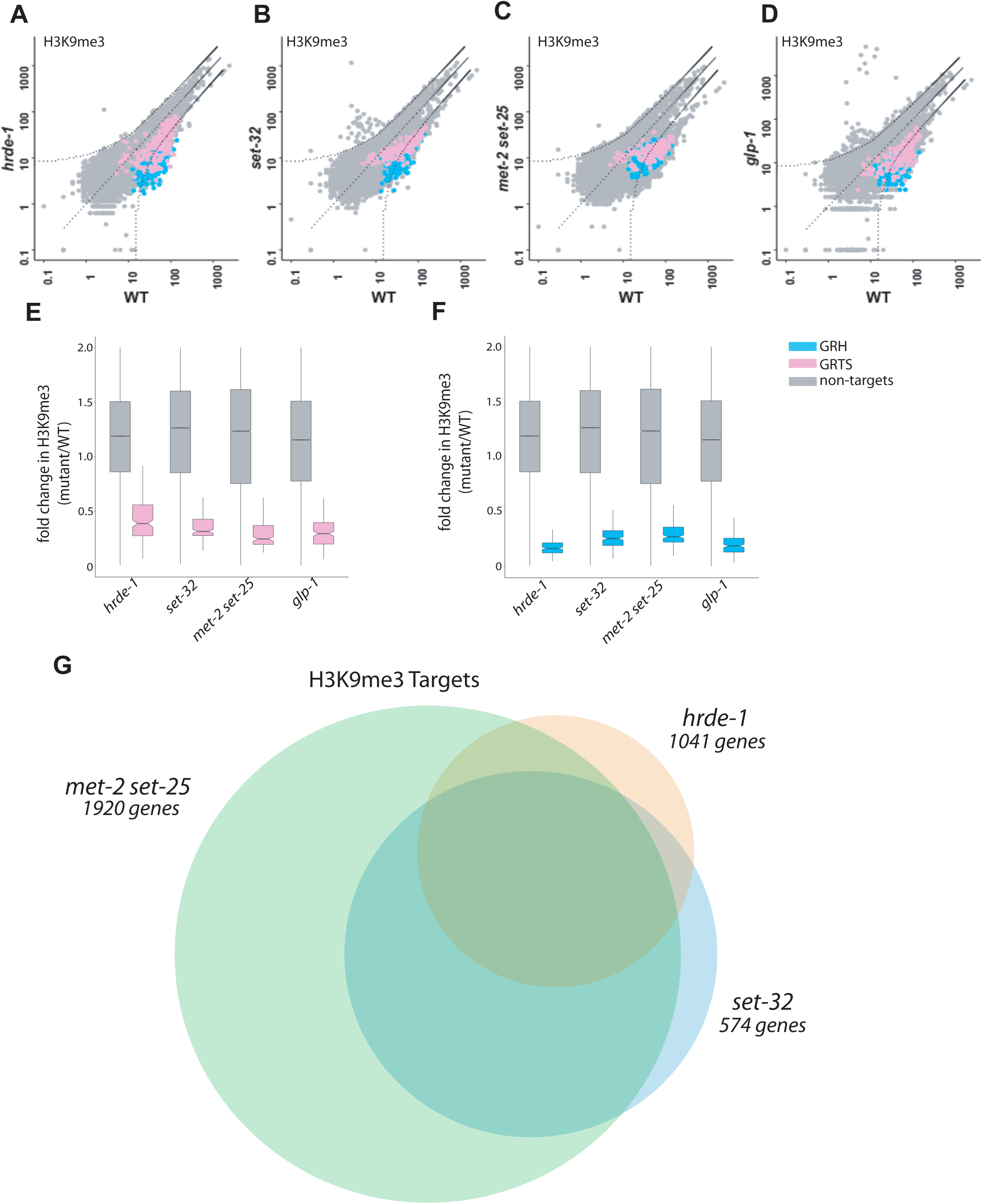
Genetic requirements for H3K9me3. **(A-D)** Scatter plots of H3K9me3 ChIP whole-genome coverage in which each point is 1kb of the genome. Mutant coverage is plotted on the y axis and WT coverage is plotted on the x axis. Curved dashed lines indicated two-fold difference calculated by FDR curves. Germline nuclear RNAi heterochromatin regions (GRH) and germline nuclear RNAi transcriptionally silenced regions (GRTS) loci are highlighted in pink and blue respectively. **(E-F)** Boxplot of whole-genome data representing fold change between mutant strains and WT. **(G)** Venn diagram of *hrde-1* (orange), *set-32* (blue), and *met-2 set-25* (green) dependent H3K9me3 genes. Dependence is defined as a 2-fold decrease in H3K9me3 signal compared to WT with a p value of <0.05 for a single gene in two replicas. Fisher’s exact test showed the following significance values for the overlapping regions: *hrde-1* and *set-32* < 2.2e-16, *hrde-1* and *met-2 set-25* < 2.2e-16, *set-32* and *met-2 set-25* <2.2e-16. (Note that boxplots and scatterplots are 1kb regions, while Venn Diagram is genes)

**Figure S8.**
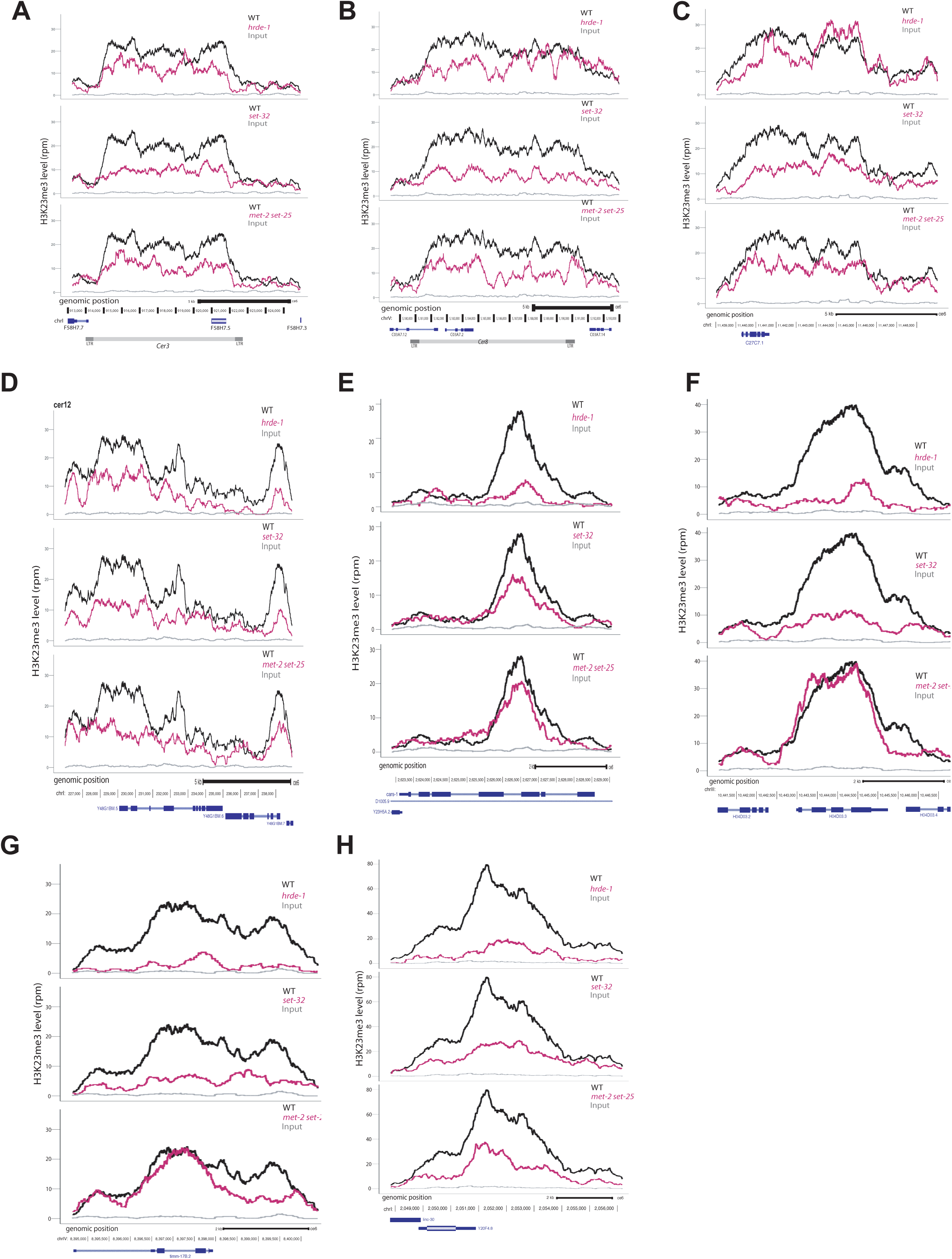
Genetic requirements for H3K23me3 at additional loci. **(A-D)** H3K23me3 levels (y axis) are plotted as a function of genomic position along the x axis in three mutant strains. Top panel: *hrde-1*, middle panel: *set-32*, bottom panel: *met-2 set-25*. For each panel WT is plotted in black, mutant is plotted in pink, grey is input in WT. The genomic loci are as follows: **(A):** *Cer3* **(B):** *Cer8* **(C):** *Cer16* **(D):** *Cer12* **(E):** *Y23H5A.7a*, a top *hrde-1*-dependent gene **(F):** *H04D03.3*, a top set-32-dependent genes **(G):** D2096.1, a top *set-32*-dependent genes **(H):** Y20F4.5, a gene depleted in all three mutant strains. All signals normalized to sequencing depth.

**Figure S9.**
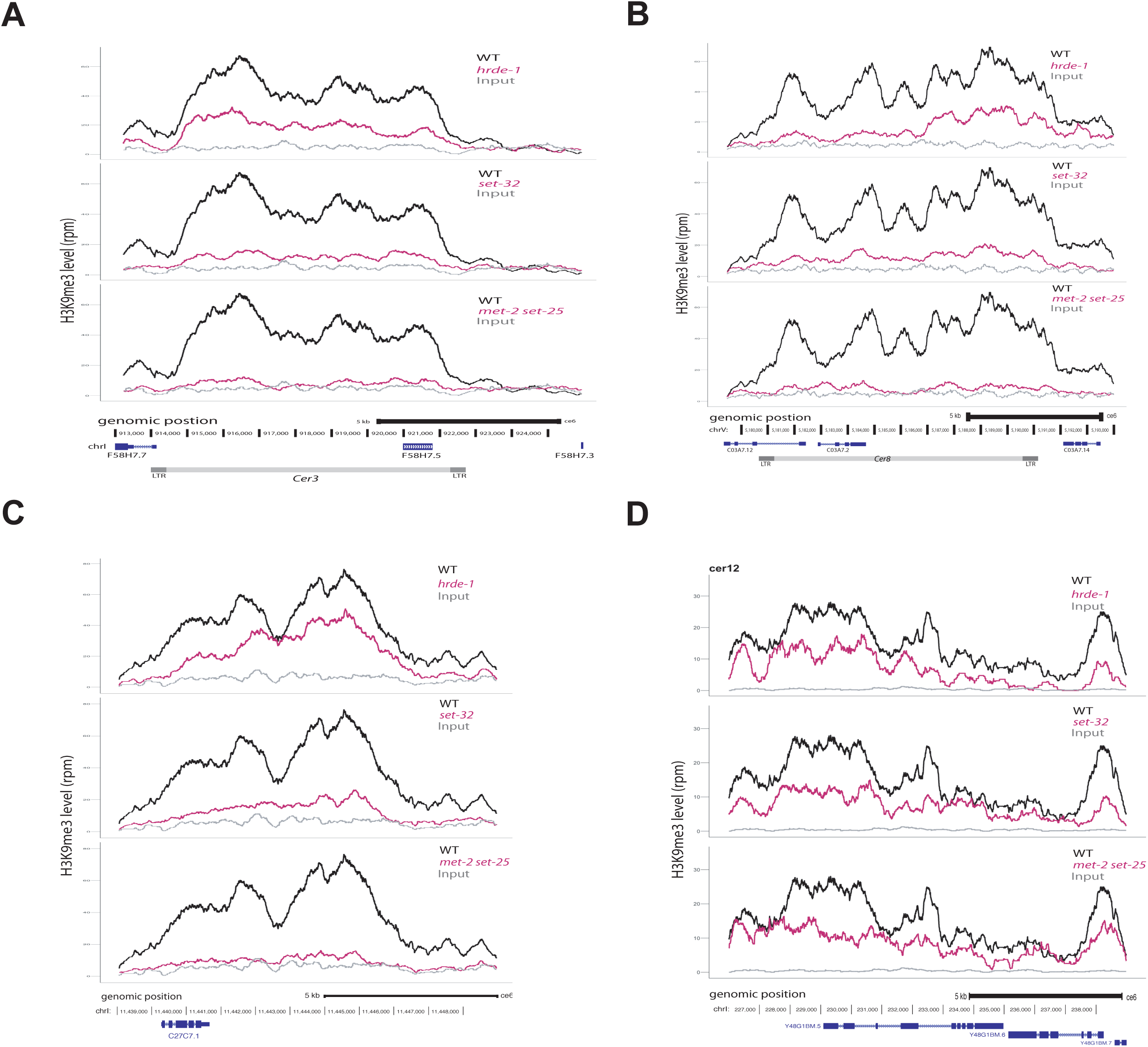
Genetic requirements for H3K9me3 at additional loci. **(A-D)** H3K9me3 levels (y axis) are plotted as a function of genomic position along the x axis in three mutant strains. Top panel: *hrde-1*, middle panel: *set-32*, bottom panel: *met-2 set-25*. For each panel WT is plotted in black, mutant is plotted in pink, grey is input in WT. The genomic loci are as follows: **(A):** *Cer3* **(B):** *Cer8* **(C):** *Cer16* **(D):** *Cer12*. All signals normalized to sequencing depth.

## Supplemental Tables

**Supplemental Table S1:**
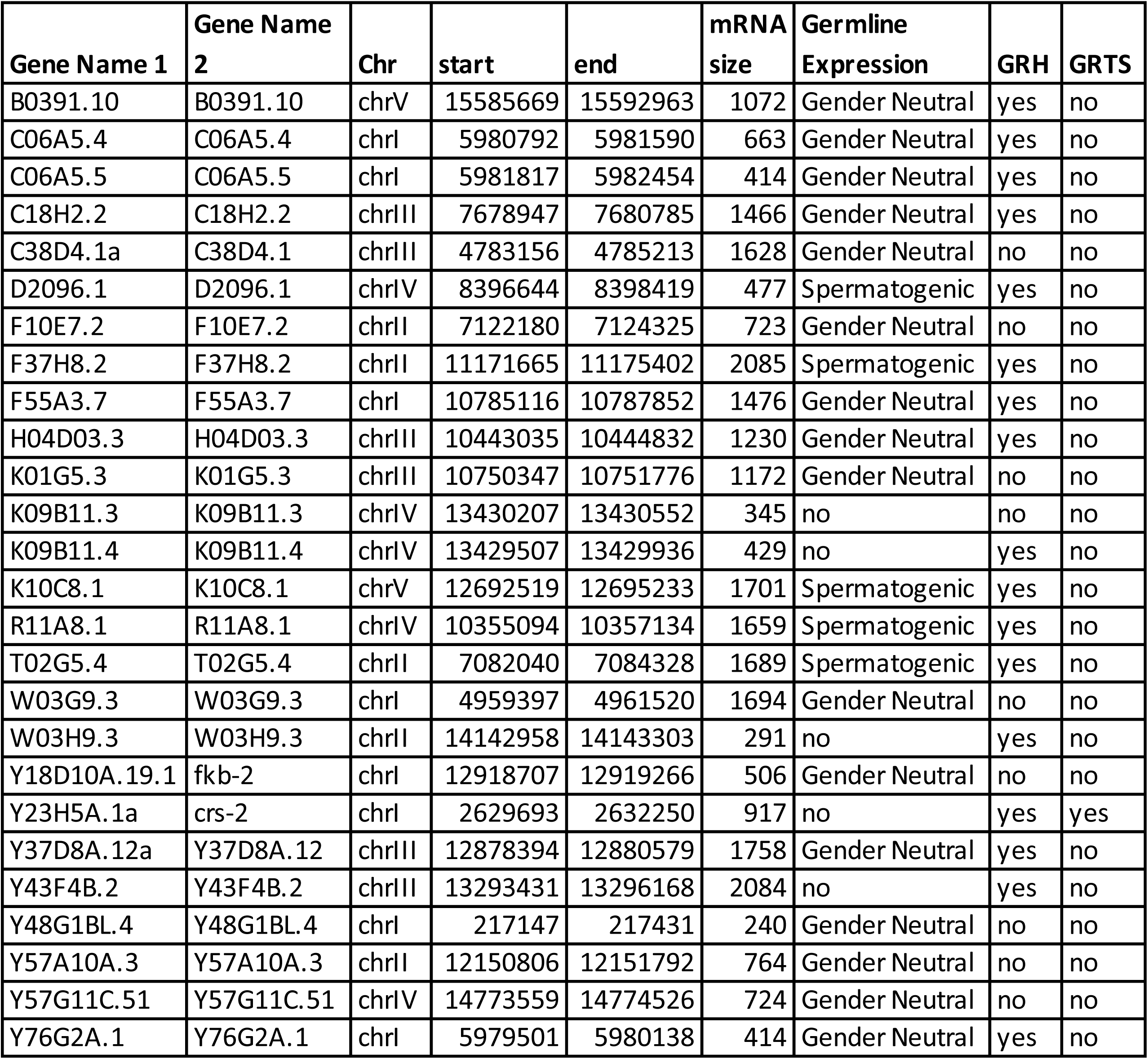
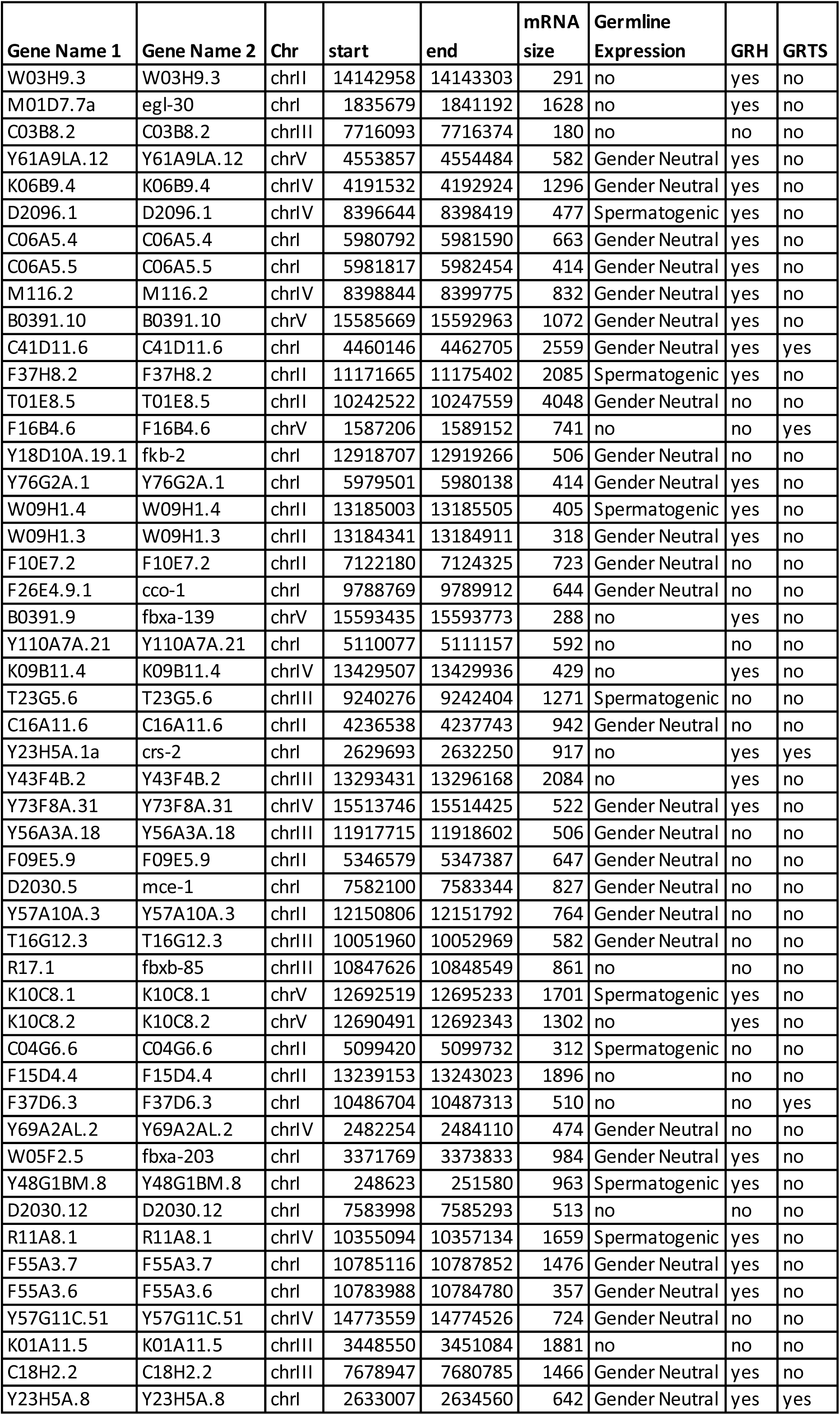
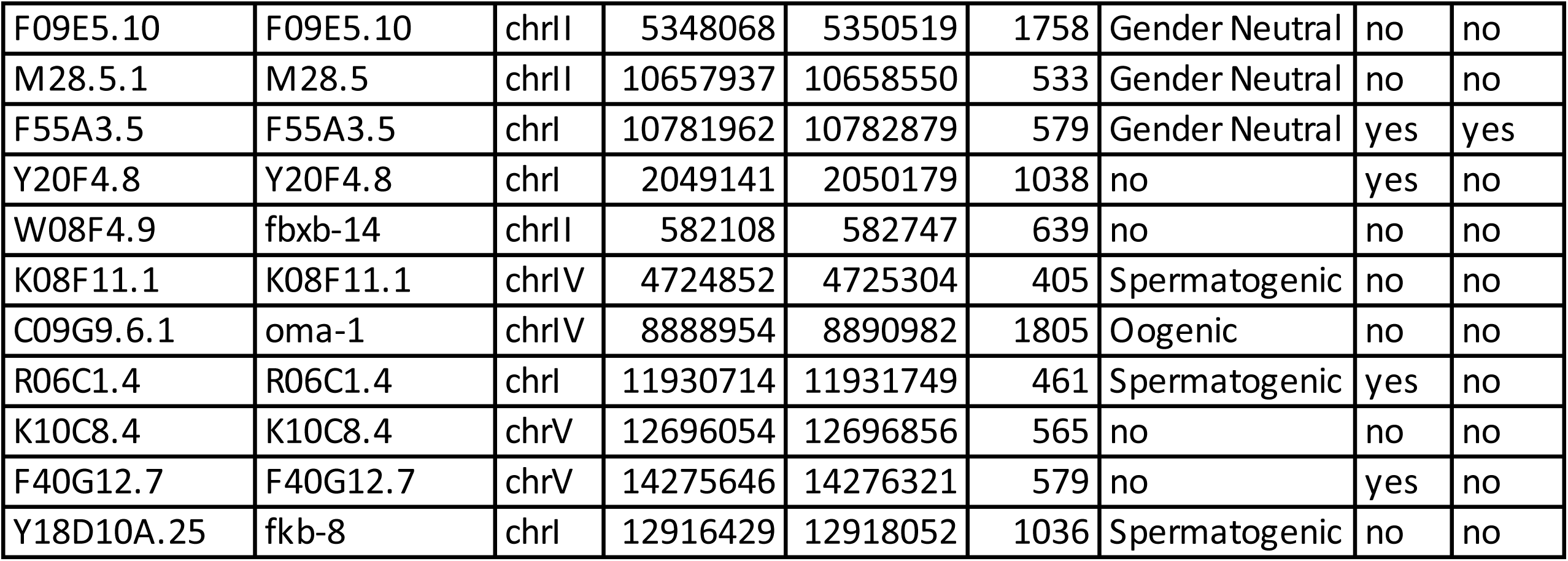
Top *set-32* and *hrde-1* dependent genes. In the first tab a list of top *set-32* dependent genes taken from annotated list of 20911 *C. elegans* genes. In the second tab a list of top *hrde-1* dependent genes. The top genes were defined as those that were 3-fold higher in WT versus mutant in two libraries with a p value of <0.05.

**Supplemental Table S2:** List of experiments, libraries, sequencing depth used in this study. ChIP-seq libraries are listed by experiment.

| Experiment | reps | library name | seq depth<br>(reads/mil) |
| --- | --- | --- | --- |
| ***oma-1* RNAi feeding, 19°C, young adult stage** |  |  |  |
| H3K23me1 ChIP-seq, WT | 1 | SG0918_lib37 | 8.5 |
| H3K23me2 ChIP-seq, WT | 1 | SG0918_lib40 | 5.2 |
| H3K23me3 ChIP-seq, WT | 1 | SG1218_lib1 | 5.5 |
| H3K23me3 ChIP-seq, WT | 2 | SG0918_lib43 | 5.5 |
| H3K23me3 ChIP-seq, set-32 | 1 | SG1218_lib3 | 4.5 |
| H3K23me3 ChIP-seq, set-32 | 2 | SG0219_lib21 | 0.9 |
| H3K23me3 ChIP-seq, met-2;set-25 | 1 | SG1218_lib4 | 5.1 |
| H3K23me3 ChIP-seq, met-2;set-25 | 2 | SG0219_lib22 | 2.8 |
| H3K23me3 ChIP-seq, hrde-1 | 1 | SG1218_lib2 | 2.8 |
| H3K9me3 ChIP-seq, WT | 1 | SG0315_lib43 | 4.3 |
| H3K9me3 ChIP-seq, set-32 | 1 | SG0615_lib32 | 3.3 |
| H3K9me3 ChIP-seq, met-2;set-25 | 1 | SG0315_lib44 | 4.2 |
| H3K9me3 ChIP-seq, hrde-1 | 1 | SG0315_lib45 | 3.5 |
| **GFP RNAi feeding, 19°C, young adult stage** | **reps** | **library name** | **seq depth** |
| H3K23me1 ChIP-seq, WT | 1 | SG0918_lib38 | 9.3 |
| H3K23me2 ChIP-seq, WT | 1 | SG0918_lib41 | 9.6 |
| H3K23me3 ChIP-seq, WT | 1 | SG0918_lib44 | 8.7 |
| H3K9me3 ChIP-seq, WT | 1 | SG0315_lib40 | 5.6 |
| ***smg-1* RNAi feeding, 19°C, young adult stage** | **reps** | **library name** | **seq depth** |
| H3K23me3 ChIP-seq, WT | 1 | SG0219_lib12 | 11.0 |
| H3K9me3 ChIP-seq, WT | 1 | SG0219_lib24 | 4.9 |
| ***oma-1* heritable, 19°C, multigenerational** | **reps** | **library name** | **seq depth** |
| *oma-1* feeding, F0, H3K23me3 ChIP, WT | 1 | SG0219_lib1 | 3.1 |
| OP50 feeding, F1 H3K23me3 ChIP, WT | 1 | SG0219_lib2 | 2.0 |
| OP50 feeding, F2 H3K23me3 ChIP, WT | 1 | SG0219_lib3 | 3.4 |
| OP50 feeding, F3 H3K23me3 ChIP, WT | 1 | SG0219_lib4 | 4.2 |
| OP50 feeding, F4 H3K23me3 ChIP, WT | 1 | SG0219_lib5 | 4.6 |
| *oma-1* feeding, F0, H3K9me3 ChIP, WT | 1 | SG1218_lib23 | 6.3 |
| OP50 feeding, F1 H3K9me3 ChIP, WT | 1 | SG1218_lib24 | 8.8 |
| OP50 feeding, F2 H3K9me3 ChIP, WT | 1 | SG1218_lib25 | 9.5 |
| OP50 feeding, F3 H3K9me3 ChIP, WT | 1 | SG1218_lib26 | 9.5 |
| OP50 feeding, F4 H3K9me3 ChIP, WT | 1 | SG0219_lib6 | 2.1 |
| **OP50 feeding, 19°C, young adult stage** | **reps** | **library name** | **seq depth** |
| H3K23me1 ChIP-seq, WT |  | SG0918_lib36 | 8.4 |
| H3K23me2 ChIP-seq, WT |  | SG0918_lib39 | 5.8 |
| H3K23me3 ChIP-seq, WT |  | SG0918_lib42 | 6.1 |
| ChIP-seq Input, WT |  | SG0918_lib45 | 8.6 |

**Supplemental Table S3:**
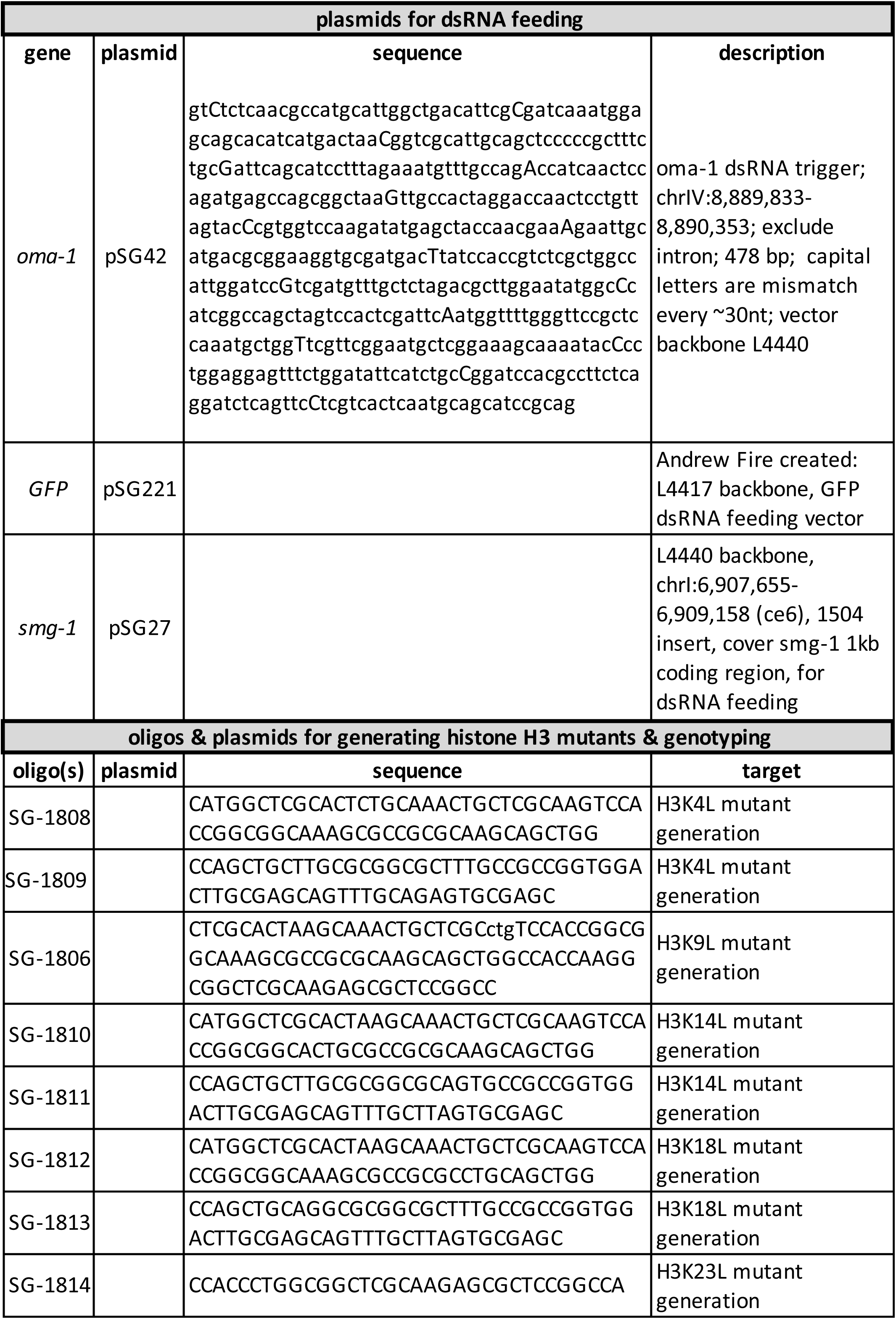
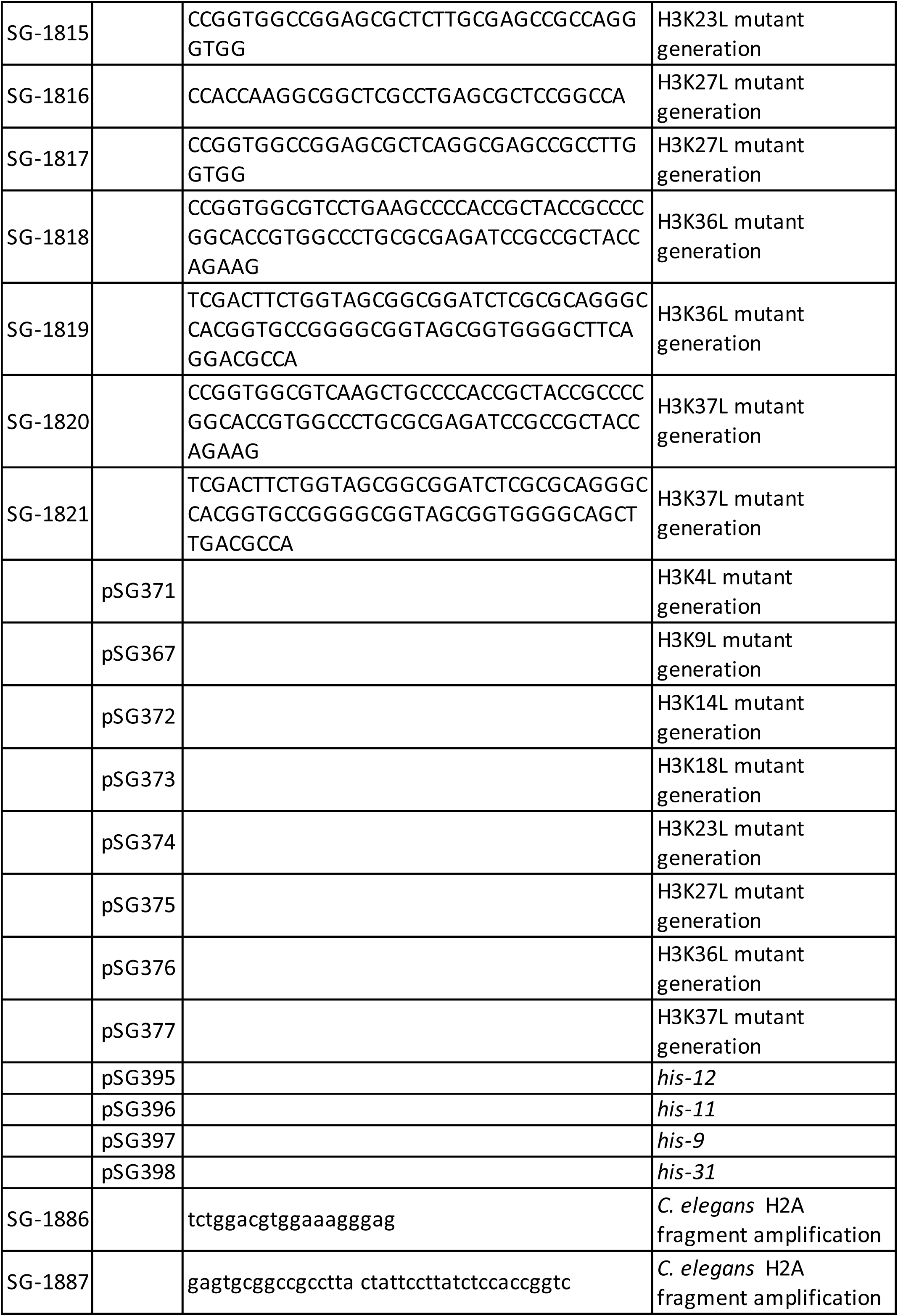
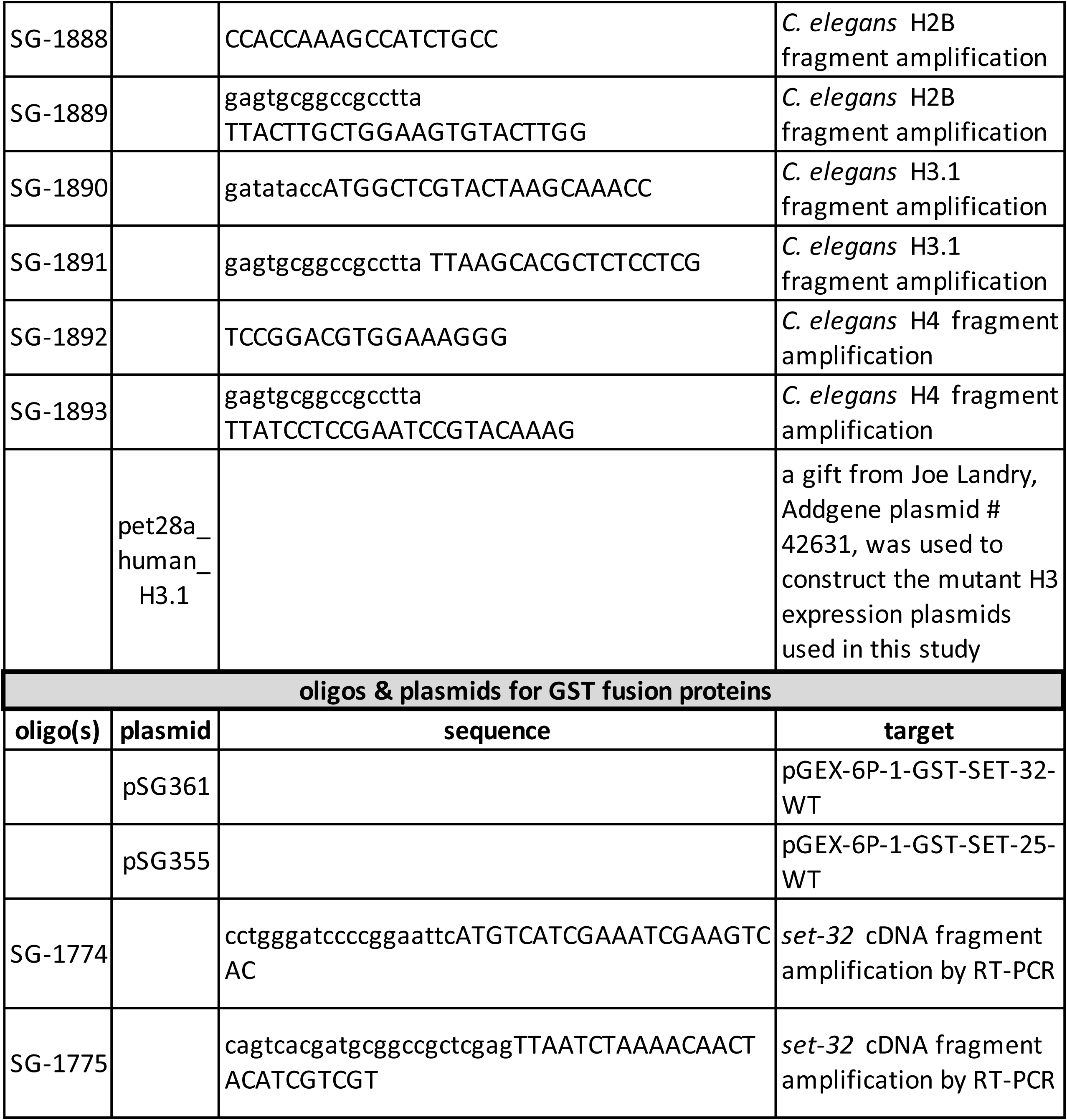
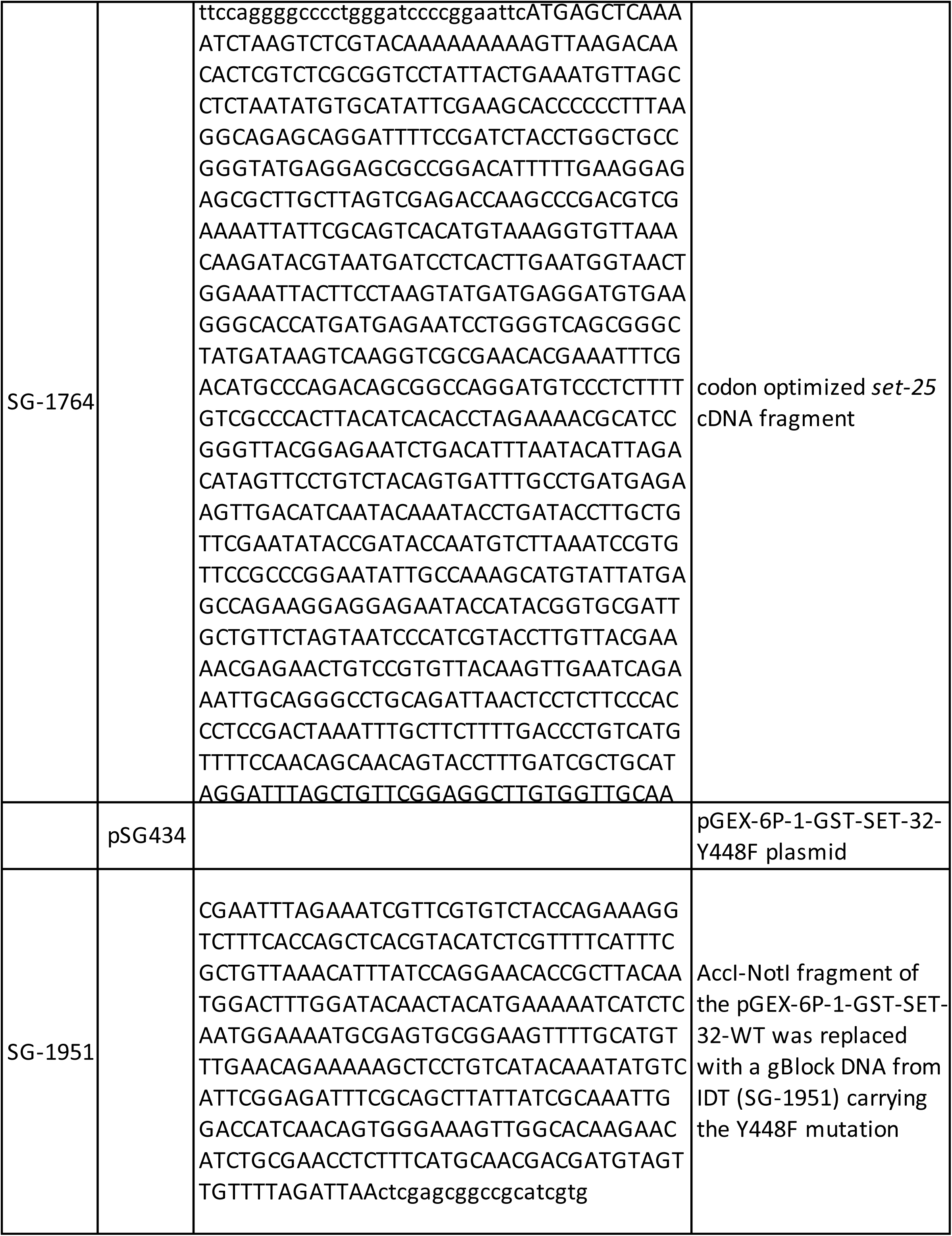
Oligonucleotides and other sequences used in this study.

